# Expansion of CD10^neg^ neutrophils and CD14^+^HLA-DR^neg/low^ monocytes driving proinflammatory responses in patients with acute myocardial infarction

**DOI:** 10.1101/2020.09.21.306118

**Authors:** Daniela Fraccarollo, Jonas Neuser, Julian Möller, Christian Riehle, Paolo Galuppo, Johann Bauersachs

**Affiliations:** Department of Cardiology and Angiology, Hannover Medical School, Hannover, Germany

## Abstract

Immature neutrophils and HLA-DR^neg/low^ monocytes expand in cancer, autoimmune diseases and viral infections, but their appearance and functional characteristics after acute myocardial infarction (AMI) remain underexplored. We found an expansion of circulating immature CD16^+^CD66b^+^CD10^neg^ neutrophils and CD14^+^HLA-DR^neg/low^ monocytes in patients with AMI, correlating with cardiac damage, function and serum levels of immune-inflammation markers. Increased frequency of immature CD10^neg^ neutrophils and elevated circulating levels of IFN-γ were linked, mainly in cytomegalovirus (CMV)-seropositive patients with high anti-CMV antibody titers and expanded CD4^+^CD28^null^ T-cells. At a mechanistic level, CD10^neg^ neutrophils enhance IFN-γ production by CD4^+^ T-cells through induction of interleukin-12. Moreover, we showed that HLA-DR^neg/low^ monocytes are not immunosuppressive but secrete high levels of pro-inflammatory cytokines after differentiation to macrophages and IFN-γ stimulation. Thus, the immunoregulatory functions of immature CD10^neg^ neutrophils play a dynamic role in mechanisms linking myeloid cell compartment dysregulation, Th1-type immune responses and inflammation in patients with AMI.

## INTRODUCTION

Despite advances in interventional therapies patients with large acute myocardial infarction (AMI) are at higher risk of heart failure morbidity and mortality.^1^ Immunity and inflammation play a key role in the pathogenesis of ischemic heart failure, and the complex role of immune cells during the wound healing process after injury is currently the focus of intensive research efforts. Understanding the immune mechanisms operating during AMI could pave the way to develop more effective strategies to prevent progressive dilative cardiac remodeling, functional deterioration and heart failure and to reduce cardiovascular adverse events. HLA-DR^neg/low^ monocytes and immature neutrophils expand in pathological conditions such as cancer, infection and inflammation,^2^ and have recently been implicated in the pathogenesis of severe COVID-19,^3–4^ but their role in immune mechanisms operating during AMI remains largely unknown.

By integrating flow cytometric immunophenotyping of monocyte, neutrophil and lymphocyte subsets, *ex vivo* experiments with sorted cells as well as bioinformatic tools this study investigated the appearance and the functional immune properties of immature neutrophils and HLA-DR^neg/low^ monocytes in patients with AMI. We also explored whether increased frequencies of immature neutrophils and HLA-DR^neg/low^ monocytes are linked to circulating levels of immune regulators and acute inflammation markers such as G-CSF, S100A9/S100A8, MMP-9, NGAL, MPO, IL-6, TNF-α, IL-1ß and IFN-γ. Using a mouse model of reperfused AMI we addressed whether immature neutrophils migrate into the ischemic myocardium.

## Methods

### Patients and study design

The study protocol is in accordance with the ethical guidelines of the 1975 declaration of Helsinki and has been approved by the local ethics committee of Hannover Medical School. Patients referred to our department for acute coronary syndrome (ACS) were included after providing written informed consent. Patients suffering from active malignant diseases or receiving immunosuppressive therapy were not included. Seventy-one patients (Table 1) were categorized into unstable angina (UA, n=11), Non-ST-elevation MI (NSTEMI, n=16), and ST-elevation MI (STEMI, n=44). Left ventricular (LV) ejection fraction was measured in 2D echocardiographic studies using bi-plane Simpson’s method. Seventeen healthy volunteers were recruited as control subjects.

**Table 1.** General Traits

|  |  | UA (n=11) | NSTEMI (n=16) | STEMI (n=44) |
| --- | --- | --- | --- | --- |
| Age (years) |  | 63.3±2.5 | 64.9±3.4 | 60.6±1.7 |
| Gender | Male/Female | 9/2 | 14/2 | 36/8 |
| BMI (kg/m<sup>2</sup>) |  | 27.5±0.9 | 28.2±0.9 | 27.5±0.7 |
| Blood analyses | LDL (mg/dL) | 92.2±19.2 | 95.5±9.5 | 138.2±7.3 |
|  | CK (IU/L) | 120.0(87.0-444.0) | 189.0(126.8-377.0) | 373.5(110.5-931.2) |
|  | CK<sub>max</sub> (IU/L) | 120.0(86.5-440.0) | 403.5(150.5-578.2) | 1343.5(574.8-1917.0) |
|  | CK-MB (IU/L) | 19.0(17.0-22.0) | 32.0(23.5-57.0) | 47.0(24.5-91.5) |
|  | Troponin (ng/L) | 12.8(5.3-22.7) | 99.0(36.7-273.5) | 337.0(84.0-962.0) |
|  | Creatinine (μmol/L) | 83.0±4.5 | 88.6±4.8 | 98.5±8.3 |
|  | CRP (mg/L) | 1.6(0.7-3.4) | 1.8(1.1-4.3) | 2.5(1.2-4.7) |

Data are presented as mean±SEM or as median (IQR). LDL, low density lipoprotein; CK, creatine kinase; CK<sub>max</sub>, maximum CK; CK-MB, creatine kinase-myocardial band; CRP, C-reactive protein.

### Flow cytometry

Venous blood was collected in EDTA tubes, stored at room temperature and processed within 1 hour of collection. White blood cell count was measured by an automated hematology analyzer (XT 2000i, Sysmex). Serum was separated within 45 minutes and stored at −80°C. For multiparameter flow cytometry whole blood (100μL) was incubated with fluorochrome-conjugated antibodies for 30 minutes at room temperature in the dark, followed by lysis of red blood cells with Versalyse Lysing Solution^®^ (Beckman Coulter).^5^ Finally, the cells were washed twice with Hanks buffer (4mL). For cell sorting by flow cytometry cells were resuspended in ice-cold FACS-staining buffer (PBS, supplemented with 0.5% bovine serum albumin and 2mM EDTA) and immunostaining was performed on ice. The following antibodies were used: anti-CD14 (Clone M5E2, 1:50 BD Biosciences); anti HLA-DR (Clone L243, 1:30 BioLegend); anti-CD16 (Clone 3G8, 1:50 BioLegend); anti-CX3CR1 (Clone 2A9-1, 1:50 BioLegend); anti-CCR2 (Clone K036C2, 1:50 BioLegend); anti-CD66 (Clone G10F5, 1:30 BioLegend); anti-CD10 (Clone HI10a, 1:20 BioLegend); anti-CD3 (Clone SK7, 1:30 BD Biosciences); anti-CD4 (Clone RPA-T4, 1:30 BD Biosciences); anti-CD28 (Clone CD28.2, 1:30 BioLegend); anti-CCR7 (Clone G043H7, 1:30 BioLegend); anti-CD45RO (Clone UCHL1, 1:30 BioLegend). Fluorescence minus one (FMO) controls were included during acquisition for gating analyses to distinguish positive from negative staining cell populations. FACS data were acquired on a GalliosTM flow cytometer and analyzed with GalliosTM software (Beckman Coulter).

### Isolation of blood mononuclear cells and neutrophils

Peripheral blood was collected in EDTA tubes and mononuclear cells (PBMC) were isolated by density gradient centrifugation using Ficoll^®^-Paque Premium (GE Healthcare Biosciences). CD14^+^HLA-DR^neg/low^/CD14^+^HLA-DR^high^ monocytes cells were FACS-sorted from PBMC. Granulocytes/neutrophils were isolated from the erythrocyte fraction by dextran sedimentation or from whole blood by immunomagnetic selection (130-104-434, MACSxpress^®^ Whole Blood Neutrophil Isolation Kit; Miltenyi Biotec), and CD10^neg^/CD10^pos^ neutrophils were separated by flow-cytometric sorting. Cells were sorted in RTL Lysis Buffer plus 1% β-mercaptoethanol (74134, RNeasy Plus Mini Kit; QIAGEN), or in sterile Sorting Medium [RPMI 1640 supplemented with 10% (v/v) Heat-Inactivated Fetal Bovine Serum (HI-FCS; A3840001; Gibco)]. Cell sorting was performed using a FACS Aria Fusion or FACS Aria IIu (BD Biosciences).

### Macrophage generation and stimulation

For in vitro differentiation of monocytes into macrophages, FACS-sorted cells were suspended at 0.5×10^6^ cells/mL in RPMI 1640 medium supplemented with 10% HI-FCS and 1% PenStrep (10378016; Gibco). CD14^+^HLA-DR^neg/low^/CD14^+^HLA-DR^high^ monocytes were cultured in 96 well plates (200μL/well) in the presence of 20 ng/mL M-CSF (216-MC-005; R&D Systems) for 4 days.^6^ Monocyte-derived macrophages [(Mb), in RPMI 1640 medium supplemented with 2% HI-FCS] were stimulated with 20 ng/mL of IFNγ [M(IFNγ), 285-IF; R&D Systems] for 48 hours.

### T-cell proliferation assays in presence of monocytes

Isolation of CD3^+^ T-cells was performed using Dynabeads^®^ Untouched™ Human T-cells Kit (11344D, Invitrogen). CD3^+^ T-cells were stained with CellTrace Violet Cell Proliferation Kit (C34571; Invitrogen) and resuspended at 1×10^6^/mL in T-Cell Activation Medium (OpTmizer™ CTS™ T-Cell Expansion culture medium supplemented with L-glutamine/PenStrep; A1048501; Gibco). CD3^+^ T-cells were co-cultured in 96 well plates with CD14^+^HLA-DR^neg/low^ and CD14^+^HLA-DR^high^ monocytes at a ratio of 1 to 1 (T-cells: monocytes). T-cells were stimulated with Dynabeads Human T-Activator CD3/CD28 (11131D; Gibco) and T-cell proliferation was assessed 4 days later by CellTrace™ Violet dilution by flow cytometry.

### T-cell activation assays in presence of CD10^neg^/CD10^pos^ neutrophils

CD4^+^ T-cells were isolated from PBMC using the MojoSort™ Human CD4 T Cell Isolation Kit (480009; BioLegend) or by flow-cytometric sorting. The CD28 MicroBead Kit (130-093-247; Miltenyi Biotec) was used for isolation of CD4^+^CD28^null^ T-cells from PBMC. CD4^+^ T-cells and CD4^+^CD28^null^ T-cells were resuspended at 1×10^6^/mL in T-Cell Activation Medium and stimulated with Dynabeads Human T-Activator CD3/CD28. For transwell experiments CD4^+^ T-cells and CD10^neg^/CD10^pos^ neutrophils were co-cultured in 24 well plates at a ratio of 1 to 2 (T-cells: neutrophils) for 24 hours. CD10^neg^/CD10^pos^ neutrophils were cultured in 0.4-μm transwell inserts (140620, Thermo Scientific™) and CD4^+^ T-cells in the well beneath the insert. In some experiments, CD10^neg^/CD10^pos^ neutrophils were cultured overnight in T-Cell Activation Medium. The cell-free supernatants derived from CD10^neg^/CD10^pos^ neutrophils were added to CD4^+^ T-cells cultured in 96-well plates (8×10^4^ cells/well) in the presence of neutralizing anti-IL-12 antibody (4μg/mL; MAB219, R&D Systems) or isotype control (4μg/mL; MAB002, R&D Systems). CD4^+^CD28^null^ T-cells were cultured with cell-free supernatants derived from CD10^neg^ neutrophils. Culture supernatants were collected after 24 hours incubation.

### LEGENDplex and ELISA assays

Blood levels of G-CSF, MMP9, S100A9/S100A8, NGAL, MPO, TNF-α, IL-6, IL-1ß and IFN-γ were measured using bead-based multiplex assays (740180; 740589; 740929; LEGENDplex™ BioLegend). Serum samples were screened for CMV-specific IgG antibodies with the CMV-IgG-ELISA PKS Medac enzyme immunoassay (115-Q-PKS; Medac Diagnostika), using a cut-off value of >0.55 AU/mL for defining seropositivity according to manufacturer’s guidelines. Levels of IFN-γ, IL-12, TNF-α, IL-6, and IL-1ß in the cell-culture supernatants were measured by ELISA (DIF50; R&D Systems) and using bead-based immunoassay (740929; LEGENDplex™ BioLegend).

### RT-quantitative PCR

RNA was isolated from cells sorted in RTL Lysis Buffer using the RNeasy Plus Mini Kit (QIAGEN) according to the manufactures’ protocol. RNA quantification and quality testing were assessed by NanoDrop 2000 (Thermo Fisher Scientific) and Bioanalyzer 2100 (Agilent). cDNA synthesis was performed using 3 ng (neutrophils) and 10 ng (monocytes) of total RNA and iScript™ Reverse Transcription Supermix (Bio-Rad). Relative quantitation of mRNA expression levels was determined with CFX96 Touch™ Real Time PCR using SsoAdvanced™ Universal SYBR Green Supermix and PrimePCR™ Primers (Bio-Rad). ß-actin (ACTB) was chosen as an endogenous control. PCR amplification was performed at initially 95 °C for 30 s followed by 40 cycles at 95 °C for 5 s and terminated by 60 °C for 30s. The delta-delta Ct method was employed for data analysis.

### Animal experiments

#### Study protocol

All animal experiments were conducted in accordance with the Guide for the Care and Use of Laboratory Animals published by the National Institutes of Health (Publication No. 85–23, revised 1985). All procedures were approved by the Regierung von Unterfranken (Würzburg, Germany; permit No. 54–2531.01-15/07) and by the Niedersächsisches Landesamt für Verbraucherschutz und Lebensmittelsicherheit (Oldenburg, Germany; permit No. 33.12-42502-04-11/0644; 33.9-42502-04-13/1124 and 33.12-42502-04-17/2702). C57Bl/6 mice of both sexes were used in this study.^7–10^

#### Mouse model of reperfused AMI

Myocardial ischemia was induced by transient left coronary artery ligation in age- and gender-matched mice. Briefly, mice were anesthetized with 2% isoflurane in a 100% oxygen mix, intubated, and ventilated using a ventilator (MINIVENT mouse ventilator model 845) with the tidal volume adjusted based on body weight (10μL/g BW). Buprenorphine (0.1 mg/kg BW) was intraperitoneally administered for postoperative pain relief. The left coronary artery was ligated with a 6-0 silk suture just below the left auricular level.^7–10^ The suture was passed through a segment of PE-10 tubing. One hour after ischemia the tube was removed to allow for reperfusion. In sham-operated control mice the ligature around the left anterior descending coronary artery was not tied.

#### Isolation of immune cells and fluorescence-activated cell sorting

Mice were anesthetized, intubated and ventilated. Blood samples were drawn from the inferior vena cava into EDTA-containing tubes. Neutrophil count was measured by an automated hematology analyzer (XT 2000i, Sysmex). After lysis of red blood cells with RBC Lysis Buffer (420301; BioLegend), cells were resuspended in ice-cold FACS-staining buffer.^7–9^ The hearts were perfused for 6 minutes with the Perfusion Buffer (113mM NaCl, 4.7mM KCl, 0.6mM KH2PO4, 0.6mM Na2HPO4), 1.2mM MgSO4, 12mM NaHCO3, 10mM KHCO3, 10mM HEPES, 30mM Taurine, 5.5mM glucose, 10mM 2,3-Butanedione monoxime), and subsequently digested for 8 minutes with the Digestion Buffer (0.2mg/mL Liberase™ Roche Diagnostics; and 400μM calcium chloride in perfusion buffer), using a modified Langendorff perfusion system. The ischemic-reperfused area and surviving myocardium were separated using a dissecting microscope. Subsequently, the heart tissue was smoothly pipetted through a sterile low waste syringe several times in order to obtain a cell suspension in Stop Buffer (perfusion buffer supplemented with 10% (v/v) HI-FCS). The cell suspension was carefully filtered through a 70μm cell strainer in a 50 mL conical tube, and the cell strainer was washed with perfusion buffer. Then, the cell supension was centrifuged at 400g for 20 minutes. The pelleted cells were washed and resuspended in ice-cold FACS-staining buffer.^7–9^ To prevent capping of antibodies on the cell surface and non-specific cell labeling all steps were performed on ice and protected from light. Cells were preincubated with Fc Block (Mouse BD Fc Block™; BD Biosciences) for 10 minutes. Subsequently, fluorochrome-conjugated antibodies were added and incubated for 30 minutes. Finally, the cells were washed twice with ice-cold FACS-staining buffer. After pre-selection in side scatter (SSC) vs. forward scatter (FSC) dot plot to exclude debris and FSC vs. Time-of-Flight (ToF) dot plot to discriminate doublets by gating single cells, blood monocytes were identified as CD45^+^/CD11b^+^/Ly6G^−^/CD115^+^ cells, blood neutrophils as CD45^+^/CD11b^+^/Ly6G^+^ cells, infarct macrophages as CD45^+^/CD11b^+^/Ly6G^−^/F4/80^+^ cells and infarct neutrophils as CD45^+^/CD11b^+^/F4/80^−^/Ly6G^+^ cells.^7–9^ The following antibodies were used: anti-CD45 (clone 104, 1:100, BioLegend/BD Biosciences); anti-F4/80 (clone BM8, 1:100, BioLegend; clone T45-2342, 1:100, BD Biosciences); anti-CD11b (clone M1/70, 1:100 eBioscience/BD Biosciences); anti-CD115 (clone AFS98, 1:100 BioLegend); anti-Ly6G (clone 1A8, 1:100 BioLegend; 1:200 BD Biosciences); anti-CD182 (clone SA044G4/clone SA045E1, 1:100 BioLegend); anti-CD101 (clone Moushi101, 1:100 eBioscience; clone 307707, 1:100 BD Biosciences). For MMP-9 and IL-1ß intracellular staining, the Cytofix/Cytoperm™ Fixation/Permeabilization Kit was used according to the manufacturer’s protocol (BD Biosciences). Antibodies included anti-MMP-9 (AF909; 1:100 R&D Systems); anti IL-1ß (ab9722; 1:100 Abcam); donkey anti-goat secondary antibody (A-11055; Invitrogen) and goat anti-rabbit secondary antibody (A-11034; Invitrogen). FMO controls were included during acquisition for gating analyses to distinguish positive from negative staining cell populations. FACS data were acquired on a Gallios™ flow cytometer and analyzed with Gallios™ software (Beckman Coulter). Cell sorting was performed using a FACS Aria Fusion (BD Biosciences). Cells were sorted in Lysis-Buffer (PreEase RNA Spin Kit, Affymetrix; PN78766, USB),^7–8^ or in sterile Sorting Medium.

#### RNA-Seq

Total RNA was isolated using PrepEase RNA Spin Kit (Affymetrix; PN78766, USB) according to the manufacturer’s instructions.^7–8^ Sorted cells were directly collected in lysis buffer and immediately processed. RNA quantification and quality testing were assessed by NanoDrop 2000 (Thermo Fisher Scientific) and Bioanalyzer 2100 (Agilent). Libraries for RNA sequencing were prepared from 30 ng total RNA; from each sample, polyA RNA was purified, converted to cDNA and linked to Illumina adapters using the Illumina TruSeq stranded mRNA Kit according to the manufacturer’s instructions. Samples were multiplexed and sequenced on an Illumina NextSeq 500 in a 75 nt single end setting using a high-output run mode. Raw BCL files were demultiplexed and converted to sample-specific FASTQ files using bcl2fastq v1.8.4 (Illumina). Residual adapter sequences present in the sequencing reads were removed with Cutadapt version 1.12. Reads (~ 40 million per sample) were aligned to the mouse reference sequence GENCODE vM8 using STAR version 2.5.2b. RNA sequencing data analysis was undertaken with the statistical programming language, R. The R package DeSeq2 (v1.14.1) was used to evaluate differential gene expression.^7–8^

### Statistical Analysis

Data are presented as mean ± SEM or as median [interquartile range] as indicated. Normality of data was assessed by Shapiro-Wilk test. Normal data were analyzed by one-way ANOVA with Tukey *post hoc* test. Mann-Whitney *U* test was used to compare two independent groups. Kruskal-Wallis test was used for comparisons of median values among three or more groups, followed by un-paired Mann-Whitney *U* test for multiple comparisons. Linear regression analysis or Spearman’s rank correlation test was used to determine relationship between variables. Values of *P*≤0.05 were considered statistically significant.

## RESULTS

### Increased circulating levels of CD14^+^HLA-DR^neg/low^ monocytes in patients with acute MI

Flow cytometric immunophenotyping was performed in whole blood from patients with unstable angina (UA) or acute MI within 24 to 72 hours of symptom onset (median 43.6 hours). A time-course analysis of monocyte subset-frequencies up to day 5 after MI is shown in Figure 1-figure supplement 1.

**Figure 1-figure supplement 1.**
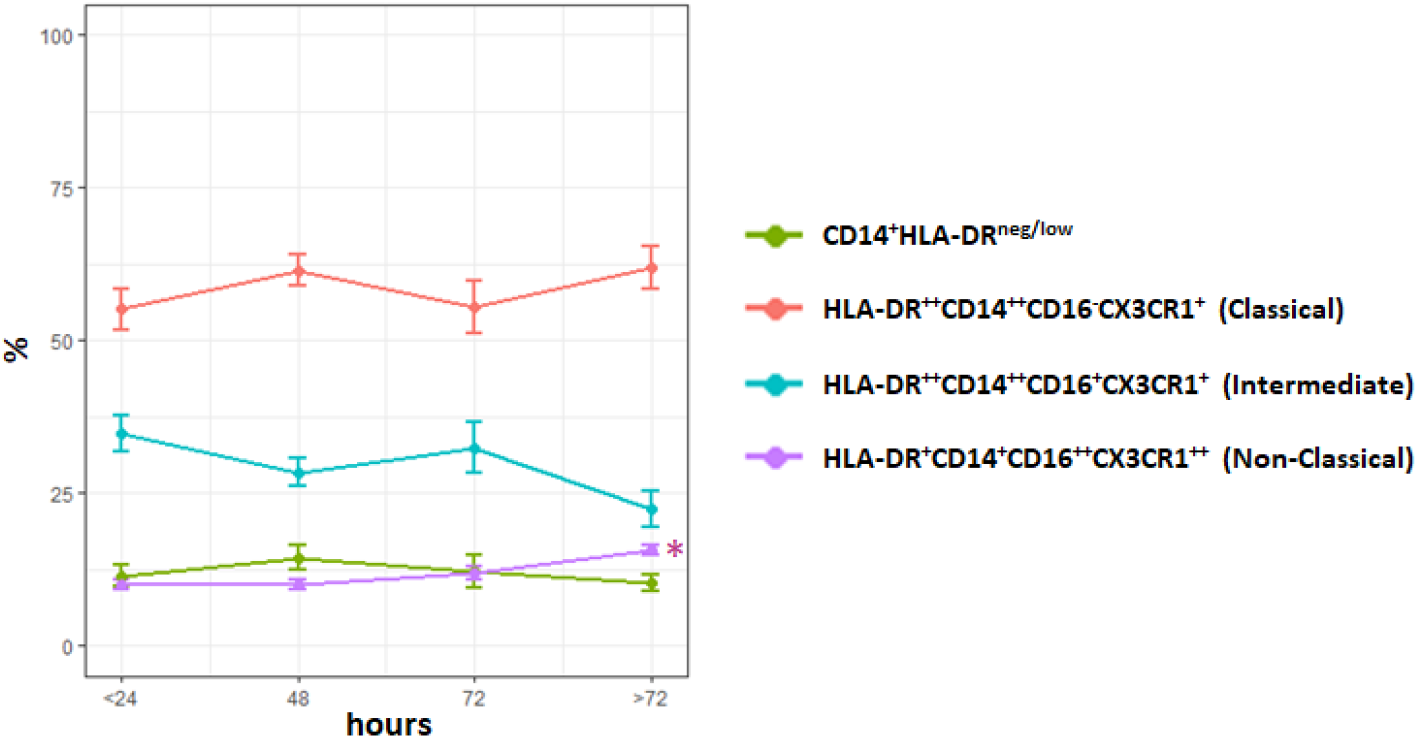
Time course analysis of monocyte subset-frequencies. Phenotypic characterization was performed within the initial 24 hours and up to day 5 after onset of symptoms in patients with ACS. \**P*<0.01 vs. ≤ 24 hours. Error bars represent SEM.

NSTEMI/STEMI patients displayed significantly higher absolute neutrophil and monocyte counts versus UA patients (Table 2). Based on HLA-DR/CD14/CD16 expression monocytes can be divided into different subsets. We detected increased circulating levels of intermediate (HLA-DR^++^CD14^++^CD16^+^CX3CR1^+^) in ACS patients versus control, and of non-classical (HLA-DR^+^CD14^+^CD16^++^CX3CR1^++^) in STEMI versus UA patients and controls (Table 2 Figure 1-figure supplement 2). There were no significant correlations between intermediate/non-classical monocytes and LV ejection fraction/CK_max_.

**Table 2.** Leukocyte Count and Monocyte Subsets

| | CTR ($n = 17$) | UA ($n=11$) | NSTEMI ($n=16$) | STEMI ($n=44$) | $P$ (K-W) |
| --- | --- | --- | --- | --- | --- |
| Neutrophil ($10^3/\mu\text{L}$) | 3.25 (2.74-3.42) | 4.05 (3.65-4.56)* | 5.72 (4.80-7.79)*† | 6.13 (5.18-7.04)*† | <0.0001 |
| Monocyte ($10^3/\mu\text{L}$) | 0.65 (0.53-0.74) | 0.74 (0.49-0.80) | 0.88 (0.72-1.03)*† | 0.99 (0.77-1.25)*† | <0.001 |
| Lymphocyte ($10^3/\mu\text{L}$) | 2.20 (1.96-2.47) | 1.82 (1.55-1.98) | 1.97 (1.76-2.49) | 2.07 (1.58-2.64) | 0.30 |
| Lymphocyte/Neutrophil ratio | 0.70 (0.57-0.79) | 0.45 (0.38-0.56)* | 0.32 (0.28-0.44)* | 0.35 (0.27-0.45)* | <0.0001 |
| Eosinophil ($10^3/\mu\text{L}$) | 0.16 (0.10-0.30) | 0.15 (0.12-0.16) | 0.20(0.14-0.34) | 0.12 (0.06-0.20) | 0.08 |
| Monocyte Classical ($n/\mu\text{L}$) | 476 (334-583) | 332 (244-388) | 510 (454-719)† | 505 (388-666)† | <0.05 |
| Monocyte Intermediate ($n/\mu\text{L}$) | 130 (73-145) | 186 (131-367)* | 204 (144-310)* | 249 (167-442)* | <0.001 |
| Monocyte Non Classical ($n/\mu\text{L}$) | 48 (30-64) | 50 (37-67) | 64 (47-108) | 102 (60-138)*† | <0.001 |

Data are presented as median (IQR). Kruskal-Wallis (K-W) test; \* $P < 0.05$ vs. Control (CTR); † $P < 0.05$ vs. UA.

**Figure 1-figure supplement 2.**
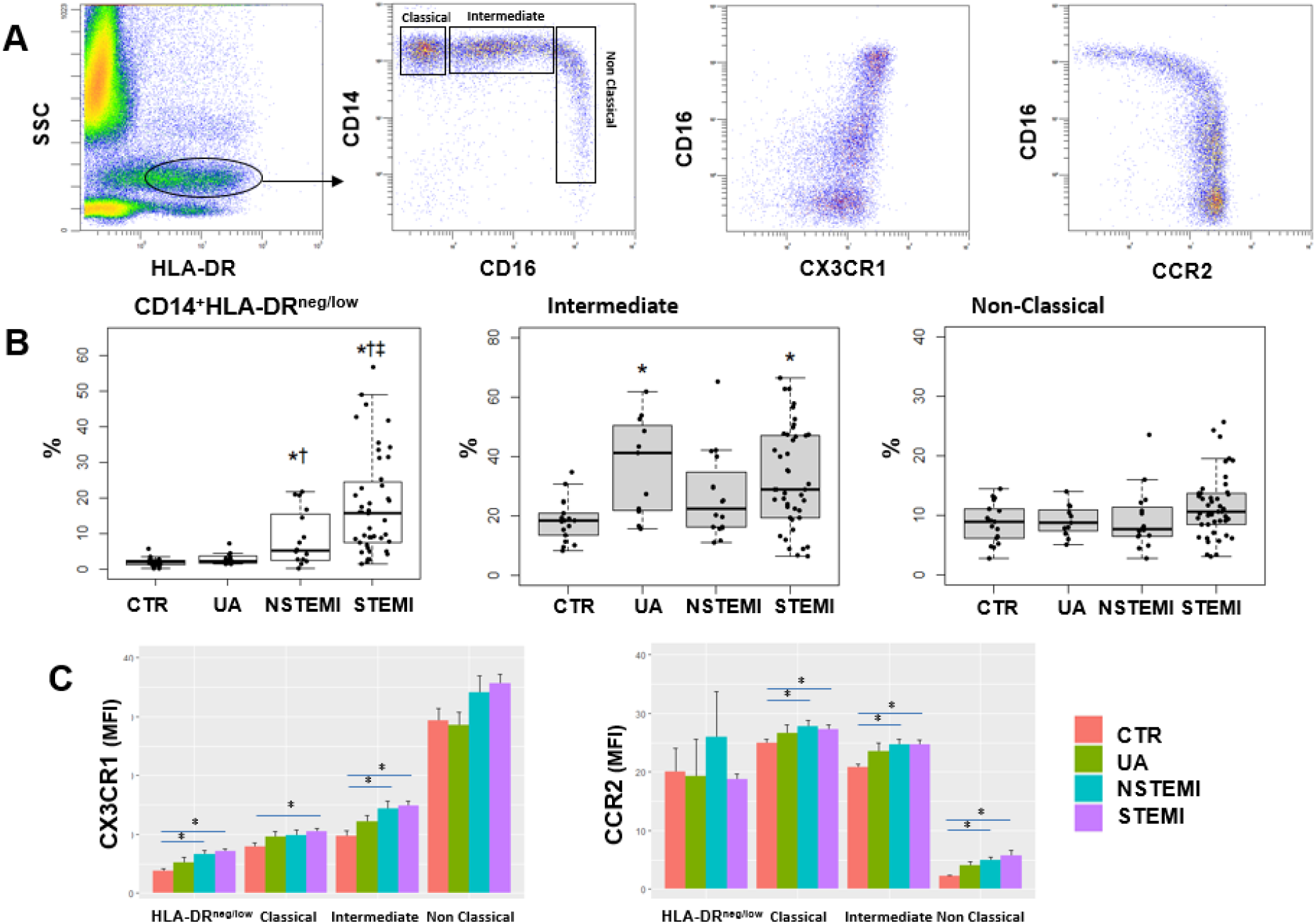
**A** Gating strategy to identify classical (HLA-DR^++^CD14^++^CD16^-^CX3CR1^+^), intermediate (HLA-DR^++^CD14^++^CD16^+^CX3CR1^+^) and non-classical (HLA-DR^+^CD14^+^CD16^++^CX3CR1^++^) monocytes. **B** Percentages of CD14^+^HLA-DR^neg/low^, intermediate and non-classical monocytes in control subjects (CTR, n=17) and in patients with unstable angina (UA; n=11), non-ST-elevation MI (NSTEMI, n=16), and ST-elevation MI (STEMI, n=44). **C** Mean fluorescence intensity (MFI) of CX3CR1 and CCR2 on monocyte subsets. \**P*<0.05 vs. CTR; †*P*<0.05 vs. UA; ‡*P*<0.05 vs. NSTEMI.

We found increased percentages and absolute numbers of circulating CD14^+^HLA-DR^neg/low^ monocytes in STEMI/NSTEMI patients as compared to UA patients (Figure 1A, 1B and Figure 1-figure supplement 2B). Linear regression analysis revealed a positive correlation between circulating levels of CD14^+^HLA-DR^neg/low^ monocytes and CK_max_ (Figure 1C) and a negative correlation with LV ejection fraction (R=0.44, *P*<0.001).

**Figure 1.**
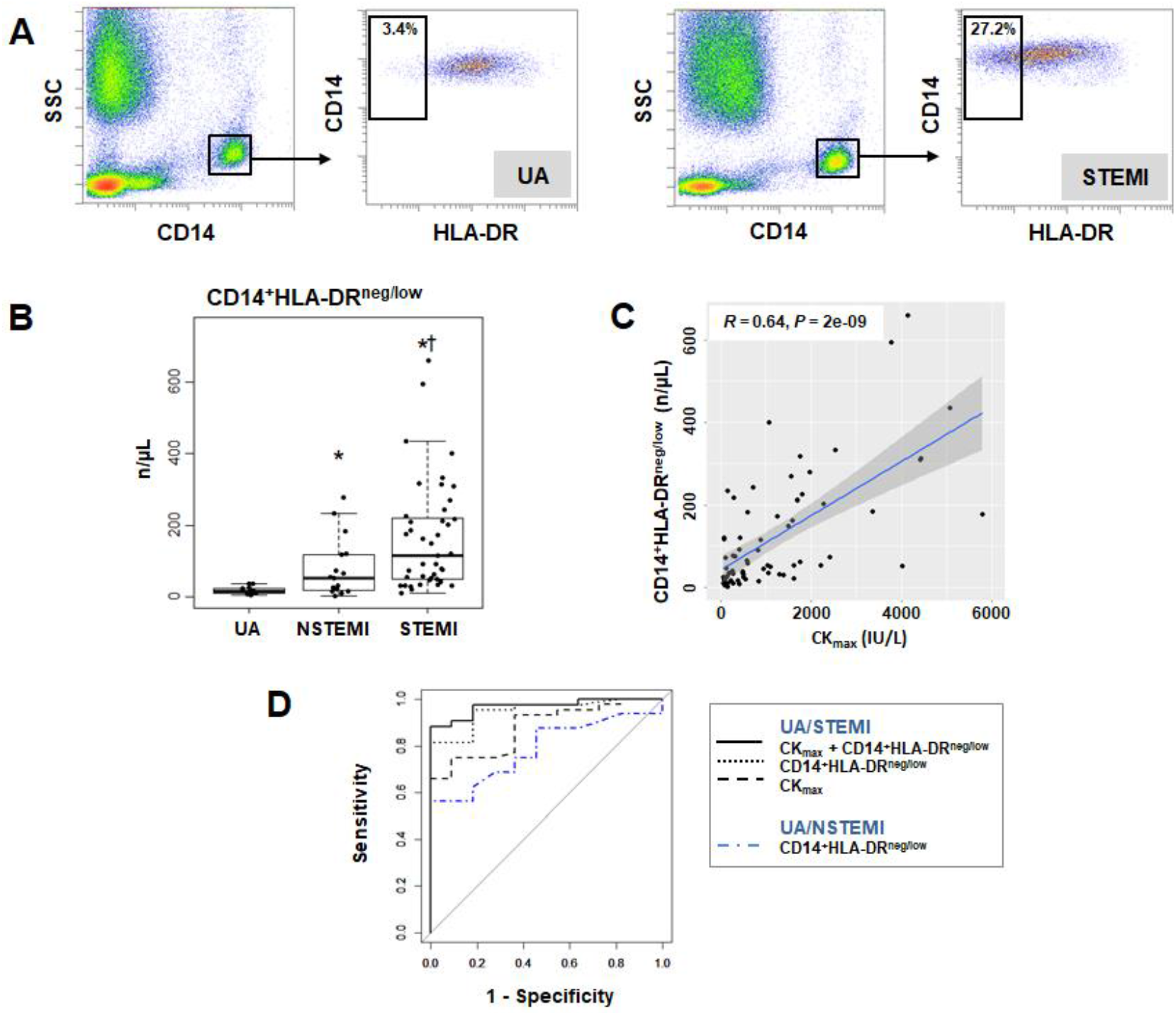
Increased circulating levels of CD14^+^HLA-DR^neg/low^ monocytes in patients with AMI. **A** Gating strategy to identify CD14^+^HLA-DR^neg/low^ monocytes. **B** Circulating levels of CD14^+^HLA-DR^neg/low^ monocytes in patients with unstable angina (UA; n=11), non-ST-elevation MI (NSTEMI, n=16), and ST-elevation MI (STEMI, n=44). **C** Linear regression analysis between circulating levels of CD14^+^HLA-DR^neg/low^ monocytes and maximum CK (CK_max_) in patients with acute coronary syndrome. **D** Receiver operator characteristic (ROC) curve of CD14^+^HLA-DR^neg/low^ monocytes discriminating UA/STEMI and NSTEMI patients and the combination of CD14^+^HLA-DR^neg/low^ monocytes (n/μL) with CK_max_. \**P*<0.05, vs. UA; †*P*<0.05, vs. NSTEMI.

Receiver operating characteristic (ROC) curve analysis based on circulating CD14^+^HLA-DR^neg/low^ monocytes (n/μL), discriminating UA and STEMI patients revealed an AUC of 0.949 (95% CI: 0.892-1; *P*<0.001) whereas a lower AUC discriminating UA and NSTEMI patients was observed (AUC=0.786; 95% CI: 0.612-0.961; *P*<0.01). By combining CD14^+^HLA-DR^neg/low^ monocytes with CK_max_ AUC was increased to 0.970; (95% CI: 0.931-1) (Figure 1D) but not in combination with LVEF (AUC=0.925; 95% CI: 0.840-1) compared to CD14^+^HLA-DR^neg/low^ monocytes alone discriminating UA and STEMI patients.

Next, we analyzed the immunoregulatory features of CD14^+^HLA-DR^neg/low^ monocytes. Using FACS-sorting, CD14^+^HLA-DR^neg/low^/CD14^+^HLA-DR^high^ cells were isolated from blood of patients with AMI (Figure 2A). Quantitative RT-PCR showed that HLA-DR^neg/low^ monocytes express high amounts of S100A9 and IL1R1 (Figure 2B). Of interest, studies in heart failure patients have provided evidence for the presence of HLA-DR^neg/low^ cells within myocardial tissue expressing high levels of S100A9.^11^

**Figure 2.**
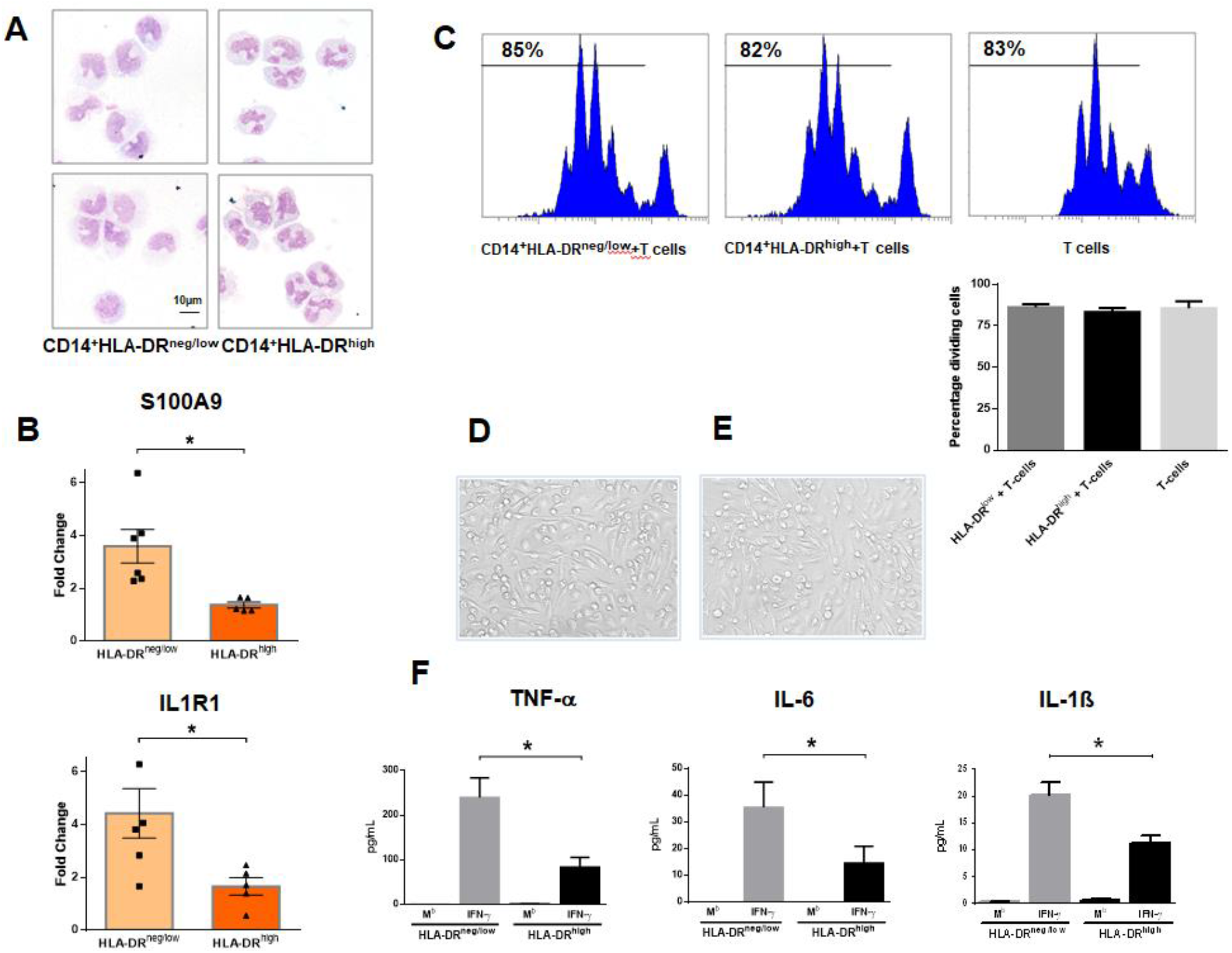
CD14^+^HLA-DR^neg/low^ monocytes from patients with AMI are not immunosuppressive but exhibit an inflammatory phenotype. **A** May-Grünwald Giemsa stained cytospin preparations of CD14^+^HLA-DR^neg/low^ and CD14^+^HLA-DR^high^ monocytes. **B** Relative RNA expression of S100A9 and IL1R1 in CD14^+^HLA-DR^neg/low^ versus CD14^+^HLA-DR^high^ monocytes. **C** T-cell proliferation in the presence of CD14^+^HLA-DR^neg/low^ or CD14^+^HLA-DR^high^ monocytes assessed by CellTrace™ Violet dilution after 96 hours of co-culture. **D** Macrophages differentiated from CD14^+^HLA-DR^neg/low^ monocytes and (**E**) CD14^+^HLA-DR^high^ cells by 4-day culture with M-CSF. **F** TNF-α, IL-6, and IL-1ß in supernatants of macrophage cultures upon stimulation with IFN-γ. M^b^=baseline. CD14^+^HLA-DR^neg/low^/CD14^+^HLA-DR^high^ cells were isolated by flow-cytometric sorting from patients with AMI (n=5-6). Data are presented as mean±SEM from independent experiments. \**P*<0.05.

CD14^+^HLA-DR^neg/low^ monocytes did not suppress T-cell proliferation (Figure 2C), indicating that the expanded population of monocytic cells in infarct patients are not immunosuppressive. Remarkably, macrophages differentiated from CD14^+^HLA-DR^neg/low^ monocytes by 4-day culture with M-CSF produced more TNF-α, IL-6, and IL-1ß upon stimulation with IFN-γ, as compared to macrophages generated from monocytes CD14^+^HLA-DR^high^ (Figure 2D through 2F). These results indicate a crucial role for CD14^+^HLA-DR^neg/low^ monocytes in the inflammatory response during AMI.

No difference was seen in the expression of CAT, CCR1, IL1R2, LCN2, MMP8, NOS2, SAAP3 and STAT3 (Figure 2-figure supplement 1A) factors dysregulated in circulating monocytes as well as in infarct macrophages in a mouse model of reperfused AMI (Figure 2-figure supplement 1B, 1C).

**Figure 2-figure supplement 1.**
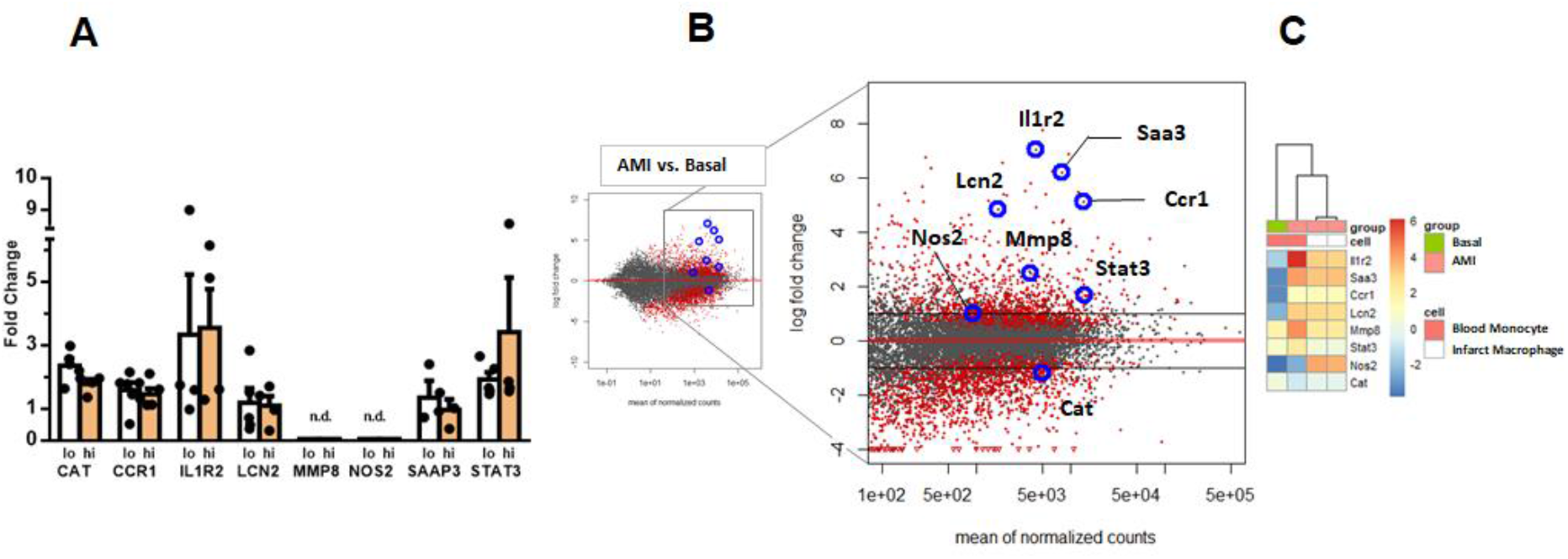
**A** RT-qPCR showing the expression of CAT, CCR1, IL1R2, LCN2, MMP8, NOS2, STAT3, SAAP3 in CD14^+^HLA-DR^neg/low^ (lo) versus CD14^+^HLA-DR^high^ (hi) monocytes FACS-sorted from blood of patients with AMI. Data are presented as mean±SEM from independent experiments. **B** MA plots showing genes regulated in circulating monocytes in a mouse model of reperfused AMI. RNA sequencing was performed on monocytes FACS-sorted from blood of sham-operated mice (Basal) and mice subjected to 1 hour of coronary occlusion followed by 6 hours of reperfusion. Genes upregulated/downregulated by AMI in monocytes were similarly regulated in (**C**) infarct macrophages FACS–sorted from the ischemic region 24 hours after AMI.

### Immature CD10^neg^ neutrophils expand in the peripheral blood from patients with acute MI

Phenotypic characterization of neutrophils was performed in whole blood. The absolute numbers and frequencies of circulating CD16^+^CD66b^+^CD10^neg^ neutrophils were significantly increased in STEMI versus UA patients (Figure 3A, 3B and Figure 3-figure supplement 1A). A time-course analysis of frequencies of CD16^+^CD66b^+^CD10^neg^ neutrophils up to day 5 after MI is shown in Figure 3-figure supplement 1B. Circulating levels of CD16^+^CD66b^+^CD10^neg^ neutrophils correlated positively with CK_max_ (Figure 3C) and negatively with LV ejection fraction (R=0.4, p<0.001).

**Figure 3.**
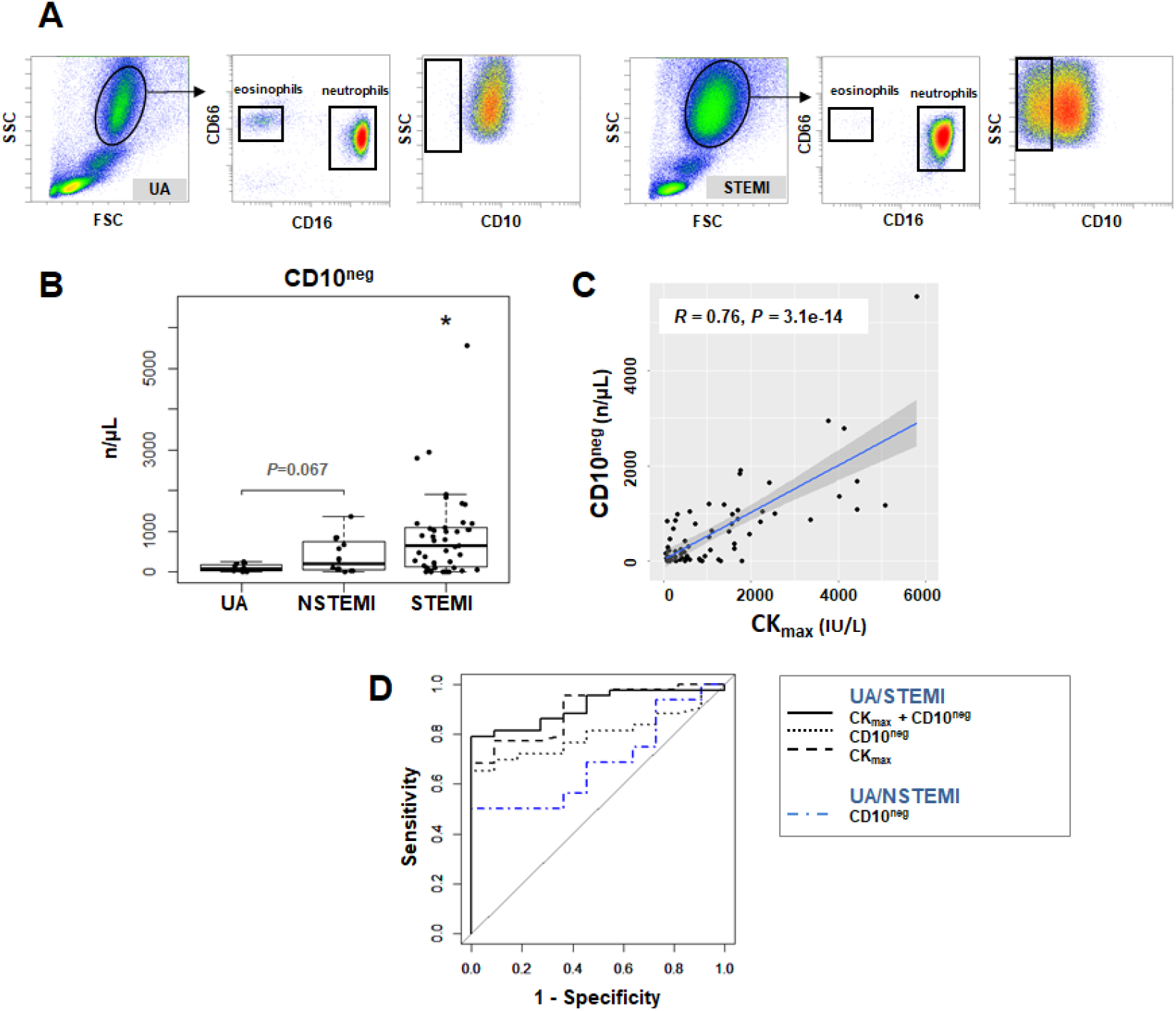
Circulating normal-density CD10^neg^ neutrophils increase in patients with AMI. **A** Gating strategy to identify CD10^neg^ neutrophils. **B** Circulating levels of CD16^+^CD66b^+^CD10^neg^ neutrophils in patients with unstable angina (UA; n=11), non-ST-elevation MI (NSTEMI, n=16), and ST-elevation MI (STEMI, n=44). **C** Linear regression analysis between circulating levels of CD10^neg^ neutrophils and maximum CK (CK_max_). **D** Receiver operator characteristic (ROC) curve of CD10^neg^ neutrophils (n/μL) discriminating UA/STEMI and NSTEMI patients and the combination of CD10^neg^ neutrophils with CK_max_ in patients with acute coronary syndrome. \**P*<0.05 vs. UA.

ROC curve analysis of circulating CD10^neg^ neutrophils (n/μL), discriminating UA and STEMI patients revealed an AUC of 0.798 (95% CI: 0.683-0.913; *P*<0.001) but a lower AUC discriminating UA and NSTEMI patients (AUC=0.687; 95% CI: 0.482-0.892; *P*=0.015). By combining CD10^neg^ neutrophils with CK_max_ or LVEF AUC was increased to 0.909; (95% CI: 0.831-0.986) and to 0.833 (95% CI: 0.691-0.974) respectively discriminating UA and STEMI patients (Figure 3D).

**Figure 3-figure supplement 1.**
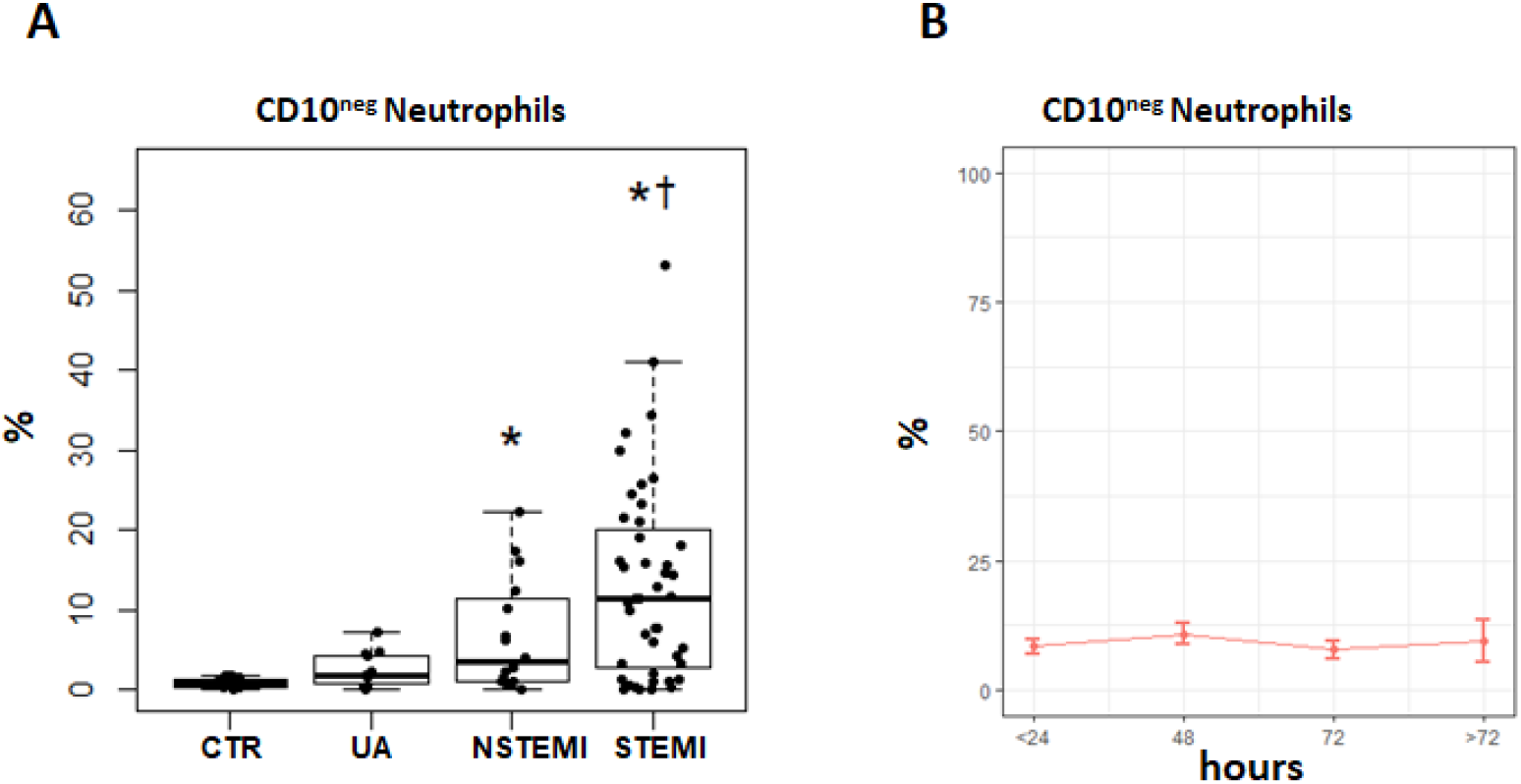
**A** Percentages of circulating CD16^+^CD66b^+^CD10^neg^ neutrophils in patients with unstable angina (UA; n=11), non-ST-elevation MI (NSTEMI, n=16), and ST-elevation MI (STEMI, n=44). **B** Time course analysis of frequencies of CD16^+^CD66b^+^CD10^neg^ neutrophils. Phenotypic characterization was performed within the initial 24 hours and up to day 5 after onset of symptoms in patients with ACS. \**P*<0.05, vs. CTR; †*P*<0.05 vs. UA. Error bars represent SEM.

CD16^+^CD66b^+^CD10^neg^ neutrophils co-purified with the erythrocyte fraction following density gradient centrifugation. Low-density neutrophils were not present in mononuclear cell fraction obtained from AMI patients. Cytospin slides were made after FACS-sorting to examine nuclear morphology (Figure 4A). We found that the majority of the CD16^+^CD66b^+^CD10^neg^ cells has an immature morphology with a lobular nucleus, while CD16^+^CD66b^+^CD10^pos^ cells are mature neutrophils with segmented nuclei (Figure 4A). These findings were obtained when neutrophils were isolated by dextran sedimentation as well as by negative selection using magnetic beads, indicating that the differences between the neutrophil subpopulations cannot be considered an artifact due to the isolation technique used.^12^ Of note, linear regression analysis revealed a strong positive correlation between the percentages of CD16^+^CD66b^+^CD10^neg^ cells and circulating levels of G-CSF (Figure 4B). AMI patients with higher systemic concentrations of G-CSF have increased CD10^neg^ neutrophils levels, suggesting G-CSF-driven immature neutrophil release/expansion.

**Figure 4.**
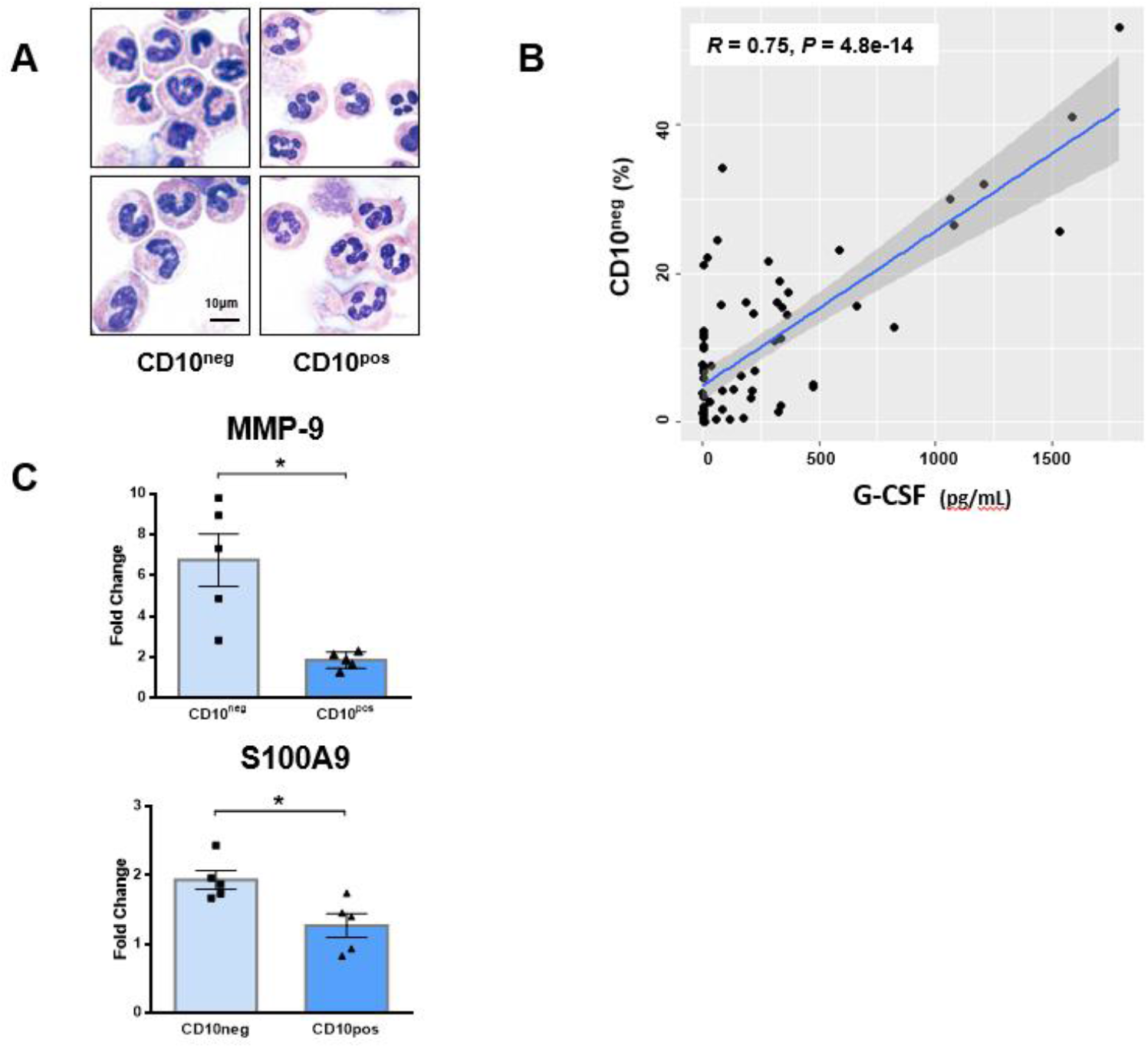
Immature CD10^neg^ neutrophils from patients with AMI express high amounts of MMP-9 and S100A9. **A** May-Grünwald Giemsa stained cytospin preparations of CD16^+^CD66b^+^CD10^neg^ (CD10^neg^) and CD16^+^CD66b^+^CD10^pos^ (CD10^pos^) neutrophils. **B** Linear regression analysis between the percentages of CD16^+^CD66b^+^CD10^neg^ neutrophils and circulating levels of G-CSF in patients with acute coronary syndrome (n=71). **C** Relative RNA expression of MMP-9 and S100A9 in CD10^neg^ versus CD10^pos^ neutrophils. CD10^neg^/CD10^pos^ neutrophils were isolated by flow-cytometric sorting from patients with AMI (n=5). Data are presented as mean±SEM from independent experiments. \**P*<0.05.

CD10^neg^ neutrophils sorted from blood of AMI patients express higher amounts of MMP-9 and S100A9 than CD10^pos^ neutrophils (Figure 4C). No difference was found in the expression of IGR1, ILR1, ILR2, MMP-8, NOS2, OLFM4 and STAT3 (Figure 4- figure supplement 1A), genes regulated in circulating neutrophils as well as in infarct neutrophils in a mouse model of reperfused AMI (Figure 4- figure supplement 1B, 1C).

**Figure 4-figure supplement 1.**
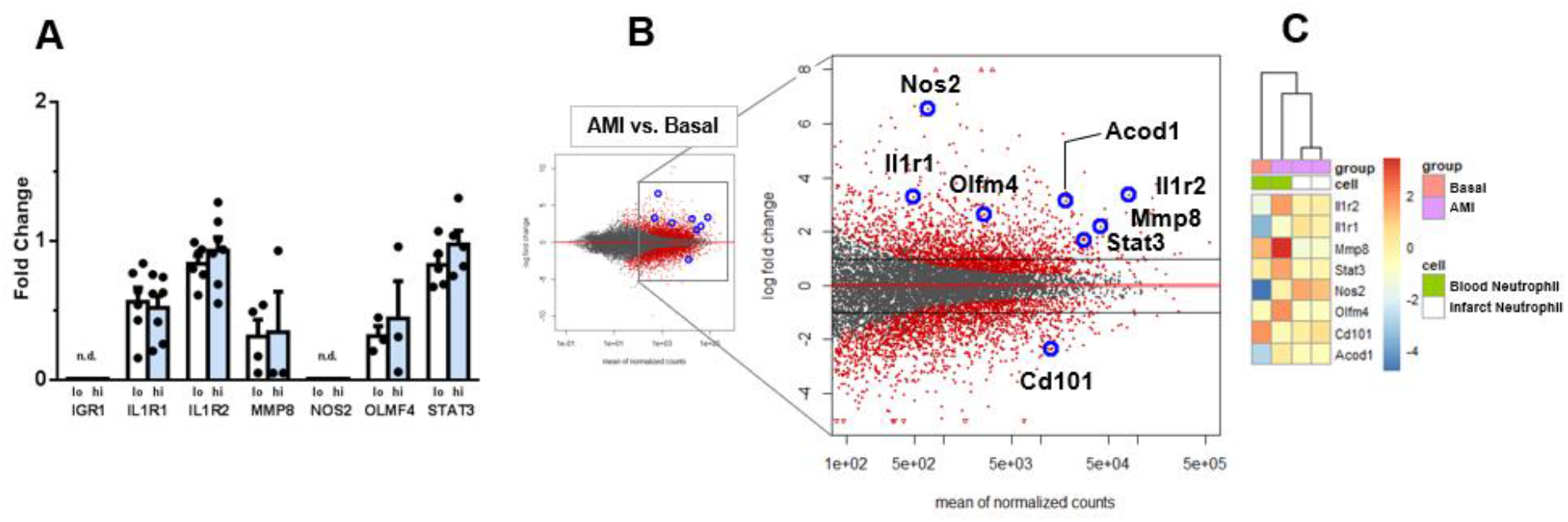
**A** RT-qPCR showing the expression of IGR1, ILR1, ILR2, MMP-8, NOS2, OLFM4, STAT3 in CD10^neg^ (lo) versus CD10^pos^ (hi) neutrophils FACS-sorted from blood of patients with AMI. Data are presented as mean±SEM from independent experiments. **B** MA plots showing genes regulated in circulating neutrophils in a mouse model of reperfused AMI. RNA sequencing was performed on neutrophils FACS-sorted from blood of sham-operated mice (Basal) and mice subjected to 1 hour of coronary occlusion followed by 6 hours of reperfusion. Genes upregulated/downregulated by AMI in neutrophils were similarly regulated in (**C**) infarct neutrophils FACS–sorted from the ischemic region 24 hours after AMI.

### Immature neutrophils are recruited to sites of cardiac injury in a mouse model of reperfused acute MI

We then investigated whether immature neutrophils had the capacity to migrate into the ischemic myocardium using a mouse model of reperfused AMI. Mouse neutrophils, unlike human granulocytes, lack CD10 expression.^13^ Using next-generation RNA sequencing we identified CD101 among the genes down-regulated by ischemia in circulating neutrophils (Figure 4-figure supplement 1B). Thus, we found that CD101 can be used as a surface marker to identify the immature neutrophil subset among the heterogeneous Ly6G^pos^Cxcr2^pos^ neutrophil populations, released into the bloodstream 90 minutes after reperfusion (Figure 5A, 5B). As revealed by morphological analysis circulating CD11b^bright^CD101^pos^ neutrophils have a mature morphology, whereas CD11b^dim^CD101^neg^ cells are immature neutrophils with ring-shaped nuclei. Of interest, a recent study in mice showed that CD101 segregates immature neutrophils from mature neutrophils during G-CSF stimulation and in the tumor setting.^14^

**Figure 5.**
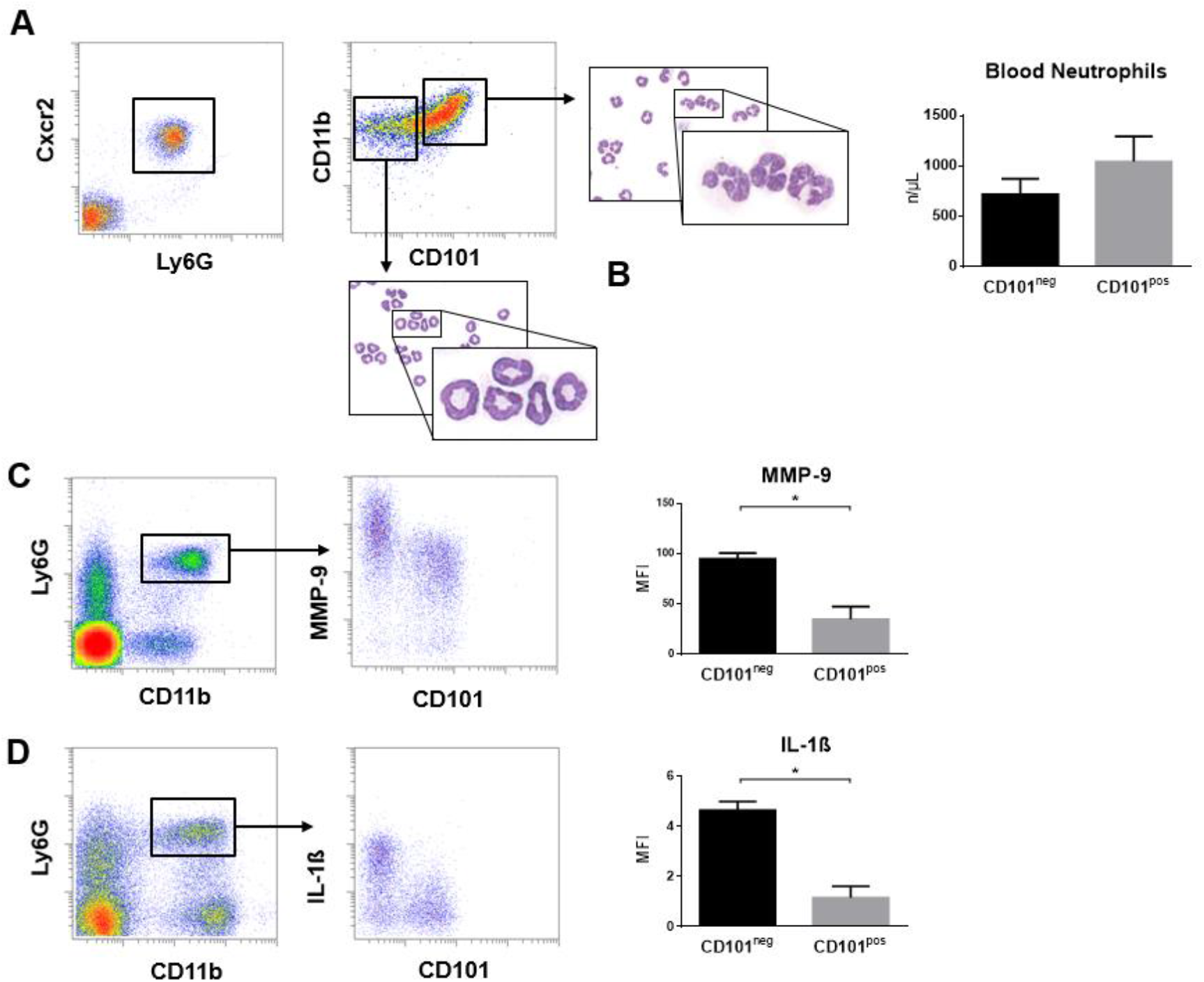
Immature CD101^neg^ neutrophils are rapidly recruited to ischemic sites and are a major source of MMP-9 and IL-1ß in the reperfused myocardium in a mouse model of AMI. **A** Representative gating strategy to identify circulating immature neutrophils among CD11b^pos^Ly6G^pos^CXCR2^pos^ cells and number of CD11b^dim^CD101^neg^ and CD11b^bright^CD101^pos^ neutrophils released into the bloodstream 90 minutes after ischemia/reperfusion. **B** May-Grünwald Giemsa stained cytospin preparations of sorted CD11b^dim^CD101^neg^ and CD11b^bright^CD101^pos^ neutrophils. **C** Flow cytometric gating strategy to identify neutrophils in the ischemic region 3 hours after reperfusion and mean fluorescent intensity (MFI) of MMP-9 on CD101^neg^ and CD101^pos^ neutrophils. **D** Flow cytometry identifying infarct neutrophils 24 hours after reperfusion and mean fluorescent intensity of IL-1ß on CD101^neg^ and CD101^pos^ neutrophils. Data are presented as mean±SEM from independent experiments (n=3-4). \**P*<0.05.

We next analyzed whether CD101^neg^ neutrophils are recruited to the injured myocardium. Preliminary experiments showed that current protocols for tissue dissociation and the recovery of neutrophils from ischemic myocardium involving long enzymatic digestion times resulted in cell activation/damage and non-specific phenotypic changes. Therefore, using a modified Langendorff perfusion system, the infarcted hearts were perfused for 6 minutes to remove blood cells and subsequently digested for only 8 minutes to preserve cell surface antigens along with expression profiles. Flow cytometry analysis of immune cells isolated from the ischemic region 3 hours after reperfusion revealed marked infiltration of CD101^neg^ neutrophils (1.26±0.19·10^5^, cells/infarct; n=4), displaying increased expression of the matrix-degrading protease MMP-9 (Figure 5C). Moreover, as shown in Figure 5D, we found that 24 hours after reperfusion CD101^neg^ neutrophils expressed IL-1ß at higher levels compared to CD101^pos^ cells. These findings suggest migration and homing of immature CD101^neg^ neutrophils to ischemic myocardium shortly after reperfusion.

### CD10^neg^ neutrophils and HLA-DR^neg/low^ monocytes are linked to levels of immune-inflammation markers

In a subgroup of patients we measured serum levels of immune inflammation markers (Table 3). MMP-9, S100A9/S100A8, NGAL, IL-6, and IL-1ß levels were higher in STEMI patients versus UA patients.

**Table 3.**
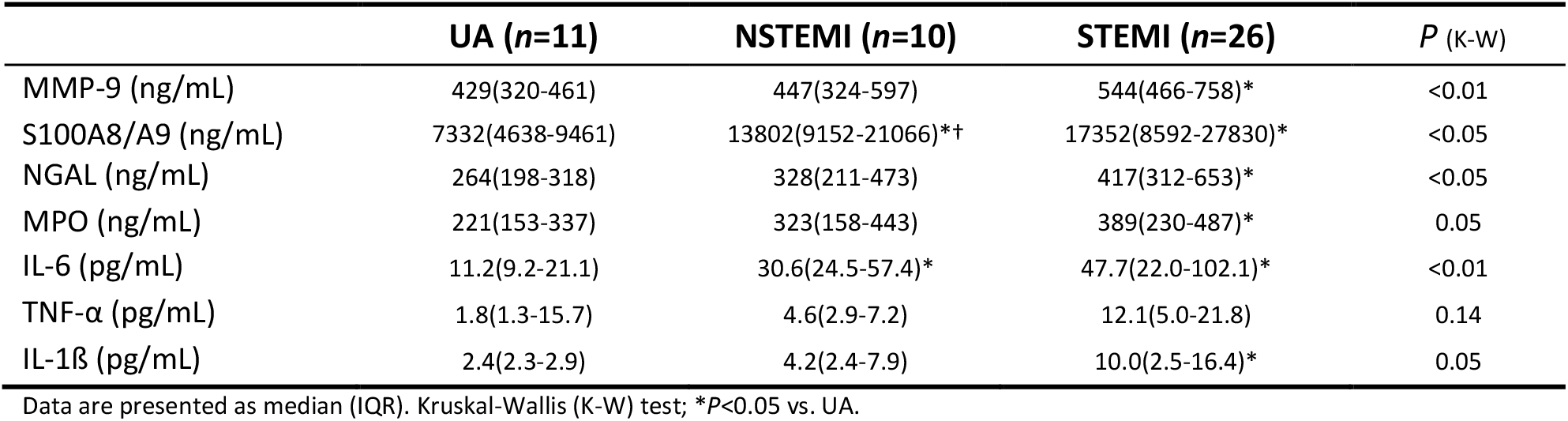
Immune Inflammation Markers

The percentages of CD14^+^HLA-DR^neg/low^ cells significantly correlated with circulating levels of MMP-9, S100A9/S100A8, IL-6, IL-1ß, TNF-α, MPO, and NGAL (Figure 6). Noticeable, CD10^neg^ neutrophils, which expand proportional to the degree of myocardial injury, significantly correlated with levels of MMP-9, S100A9/S100A8, NGAL, MPO, IL-6, and IL-1ß (Figure 6).

**Figure 6.**
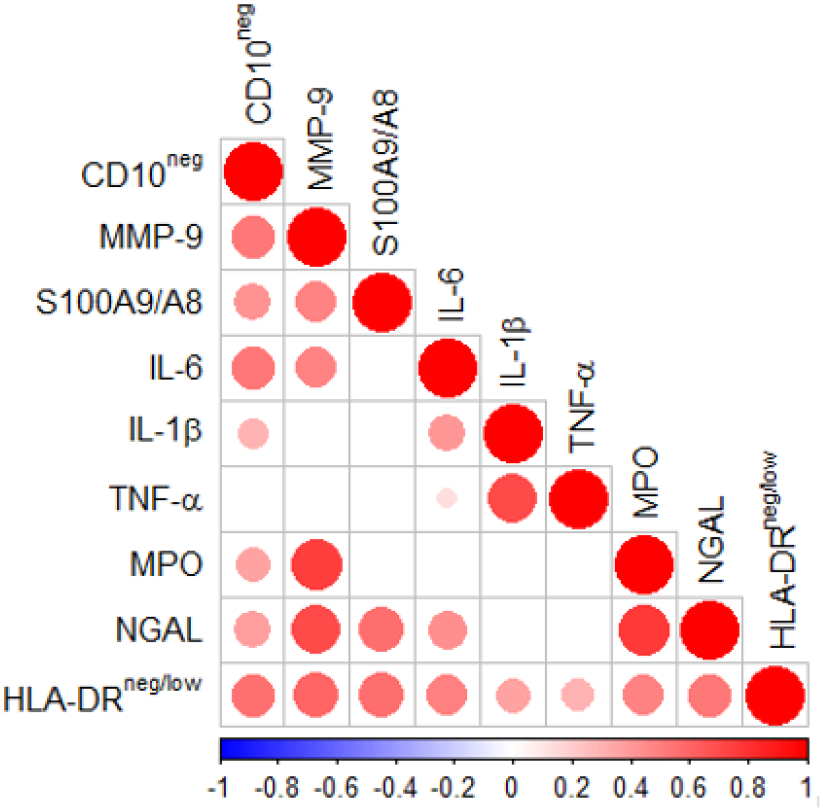
Spearman-correlation matrix of CD16^+^CD66b^+^CD10^neg^ neutrophils (%), CD14^+^HLA-DR^neg/low^ monocytes (%) and circulating levels of MMP-9, S100A9/S100A8, IL-6, IL-1ß, TNF-α, MPO, and NGAL (levels). Each circle illustrates a significant correlation between different couples of parameters (*P*<0.05). The correlation coefficient is colored and sized up according to the value; square leaved blank indicates not significant correlation.

### Elevated circulating levels of IFN-γ in cytomegalovirus-seropositive patients with expanded CD10^neg^ neutrophils and increased frequency of CD4^+^CD28^null^ T-cells

A crucial role for neutrophils in the orchestration of adaptive immunity is emerging.^15–16^ To investigate the potential immunoregulatory properties of immature CD10^neg^ neutrophils we performed flow cytometric immunophenotyping of CD4^+^ T-cells and investigated circulating levels of IFN-γ in a subgroup of patients. Contrary to some reports, circulating levels of naive (CCR7^+^CD45RA^+^), central memory (CCR7^+^CD45RA^-^), effector memory (CCR7^-^CD45RA^-^), terminally differentiated effector cells (EMRA, CCR7^-^CD45RA^+^) and CD4^+^CD28^null^ T-cells were not significantly different among patients with ACS (Table 4).

**Table 4.**
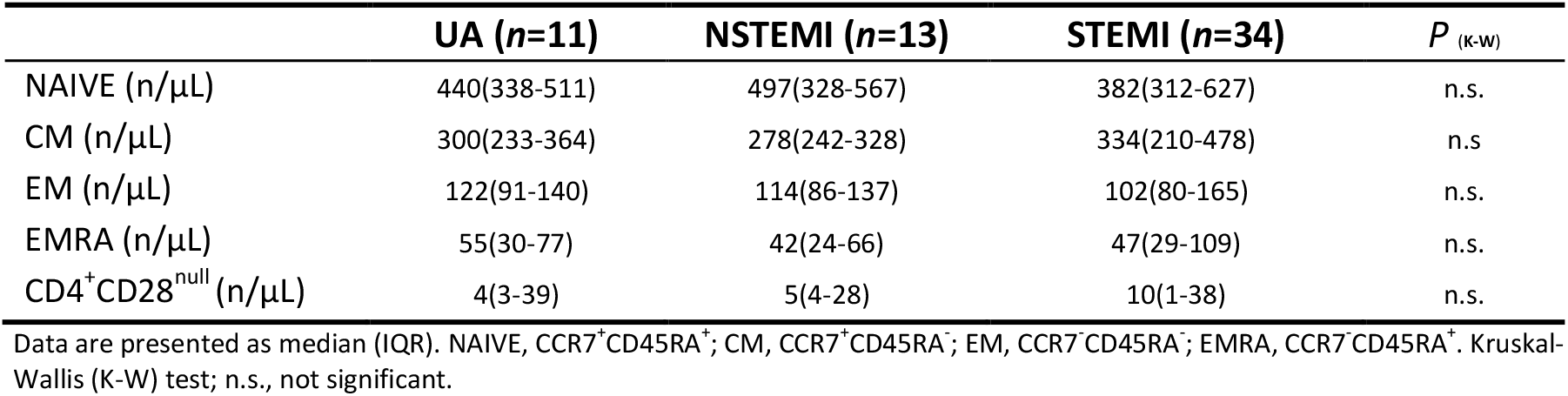
CD4^+^ T-cells Subsets

Altered T-cell homeostasis and increased frequencies of circulating CD28^null^ T-cells have been linked to cytomegalovirus (CMV) seropositivity.^17–19^ To study the impact of CMV on CD4^+^CD28^null^ T-cells frequency, UA, NSTEMI and STEMI patients were stratified according to CMV serostatus. We found that CD4^+^ T-cells lacking the costimulatory molecule CD28 showed expansion across all CMV-seropositive (CMV^+^) patients (Figure 7A, 7B). Frequency distribution of CD4^+^CD28^null^ T-cells (log_10_ transformed to improve visualization) appeared bimodal and analyzing separately in CMV^+^ and CMV-seronegative (CMV^-^) patients, the median was significantly higher by a factor 14.1 in CMV^+^ patients. (Figure 7-figure supplement 1A). Moreover, CD4^+^CD28^null^ frequency positively correlated with CMV-IgG antibody levels (R=0.6, *P*<10^−5^). Therefore, we believe that the expansion of CD4^+^CD28^null^ T-cells may not be a direct result of coronary events but appears to be related to the CMV-induced immune changes secondary to repeated antigen exposure.

**Figure 7.**
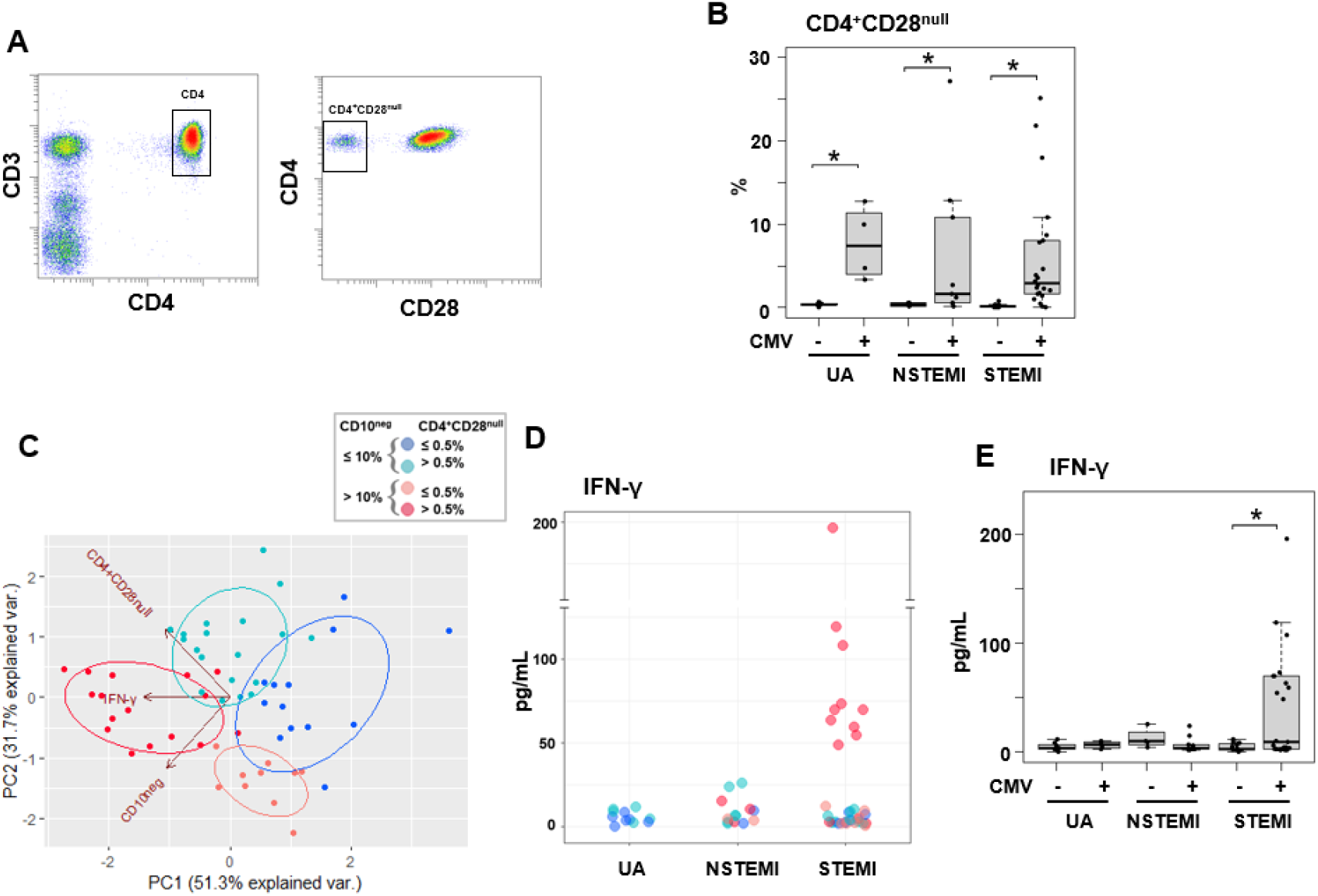
Elevated IFN-γ levels in cytomegalovirus-seropositive patients with expanded CD10^neg^ neutrophils and increased frequency of CD4^+^CD28^null^ T-cells. **A** Gating strategy indentifying CD4^+^CD28^null^ T-cells. **B** Frequency of CD4^+^CD28^null^ T-cells in patients with acute coronary syndrome stratified according to cytomegalovirus (CMV) serostatus. **C** Principal component analysis (PCA) showing clustering according to circulating levels of IFN-γ, CD10^neg^ neutrophils and CD4^+^CD28^null^ T-cells and **D** scatter plot showing IFN-γ levels according to frequency of CD10^neg^ neutrophils and CD4^+^CD28^null^ T-cells. Patients were stratified based on frequency of CD10^neg^ neutrophils (<10% or >10%) and frequency of CD4^+^CD28^null^ T-cells (<0.5% or >0.5%). **E** circulating IFN-γ levels stratified according to CMV serostatus. UA (n=11), NSTEMI (n=13), and STEMI (n=34). \**P*≤0.05.

We defined expansion of CD4^+^CD28^null^ T-cells frequencies a non-parametric, upper outlier limit (upper quartile+1.5×interquartile range) as previously reported.^17^ The subgroup of CMV^-^ patients with unstable angina was considered as reference group. (Figure 7-figure supplement 1B). Similarly we derived a cut-off for expansion of CD10^neg^ neutrophils but taking as reference group the whole cohort of patients with UA since frequency expansion appeared prevalently due to the grade of coronary disease, not to CMV-seropositivity and both contributions to cell expansion could not be dissected (Figure 7-figure supplement 1C, 1D) Then, in order to highlight relationship among IFN-γ levels, CD10^neg^ neutrophils, CD4^+^ T-cell subsets and CMV-seropositivity we performed hierarchical clustering stratifying patients with ACS according to the expansion cut-offs above described. This individuated 4 subgroups of patients with frequency of CD10^neg^ neutrophils (≤10% or>10%) and frequency of CD4^+^CD28^null^ T-cells (≤0.5% or >0.5%). In the derived heatmap IFN-γ, CD10^neg^ neutrophils, CD4^+^CD28^null^, EMRA, and EM CD4^+^ T-cells were grouped together showing similar patterns (Figure 7-figure supplement 1E), indicating that persistent CMV infection is associated with expansion of the effector memory CD4^+^ T-cell compartment and higher IFN-γ levels in patients with increased frequency of CD10^neg^ neutrophils.

**Figure 7-figure supplement 1.**
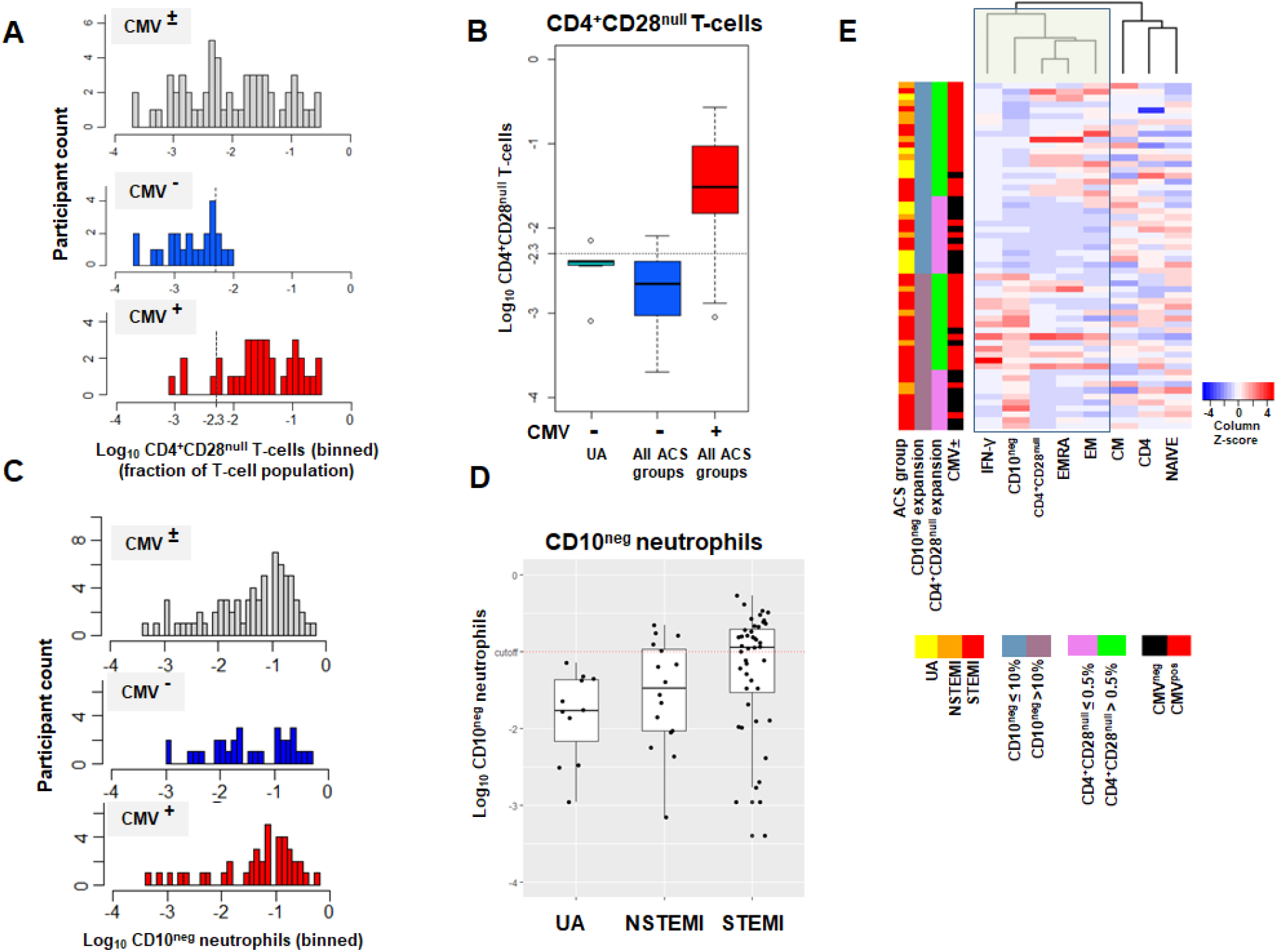
**A** CD4^+^CD28^null^ T-cell frequency distribution (log_10_ -transformed CD4^+^ T-cell fractions) of CMV± (top, n=58), CMV^-^ (middle, n=23) and CMV^+^ (bottom, n=35) acute coronary syndrome (ACS) patients. CD4^+^CD28^null^ T-cells displayed a bimodal distribution related to CMV-seropositivity. **B** Boxplots show the log_10_ -transformed frequencies of CD4^+^CD28^null^ T-cells in CMV^-^-UA, CMV^-^ (blue) and CMV^+^ (red) -ACS patients. Expansion index (dotted line) was calculated as UQ+1.5xIQR of CMV^-^-UA patients chosen as reference group. CD4^+^CD28^null^ T-cell frequency more than 0.5% was considered as an index of expansion; UQ (upper quantile), IQR (inter-quantile range). **C** CD10^neg^ neutrophils (CD10^neg^) frequency distribution (log_10_-transformed) of CMV± (top, n=71), CMV^-^ (middle, n=31), and CMV^+^ (bottom, n=40) ACS patients. **D** Boxplots show the log_10_ -transformed frequencies of CD10^neg^ in UA, NSTEMI and STEMI patients. Expansion index (dotted line) was calculated as UQ+1.5xIQR of UA patients. According, patients with CD10^neg^ frequency more than 10% had expansion. **E** Scaled frequencies of CD4^+^CD28^null^ T-cells and CD10^neg^ neutrophils stratified by criteria of cell expansion. Hierarchical clustering performed on columns highlights the relationship among CD10^neg^ neutrophils, CD4^+^CD28^null^ T-cells, IFN-γ production and CMV seropositivity.

To better highlight the relationship among IFN-γ, CD10^neg^ neutrophils and CD4^+^CD28^null^ T-cells we also performed principal component analysis (PCA) that showed clustering according to elevated circulating levels of IFN-γ, high levels of CD10^neg^ neutrophils and peripheral expansion of CD4^+^CD28^null^ T-cells (Figure 7C). The highest IFN-γ levels were found in STEMI patients with expanded CD10^neg^ neutrophils (>10%) and increased frequency of CD4^+^CD28^null^ T-cells (Figure 7D). Not surprisingly, when stratified according to CMV serostatus, maximum levels of circulating IFN-γ among ACS patients were detected in CMV-seropositive STEMI patients displaying increased levels of CD10^neg^ neutrophils (Figure 7E), indicating a relation among expansion of immature CD10^neg^ neutrophils, CMV seropositivity and strongly enhanced levels of IFN-γ in patients with large AMI.

### CD10^neg^ neutrophils via induction of interleukin-12 enhance priming for IFN-γ production by CD4^+^ T-cells

Environmental factors such as CMV infection can induce changes in CD4^+^ T-cell phenotype and function. Consequently, to provide a mechanistic understanding of the cellular basis for raised IFN-γ in CMV-seropositive patients with expanded CD10^neg^ neutrophils, we investigated IFN-γ secretion by CD4^+^ T-cells isolated from CMV^-^/CMV^+^ patients and its potential link to interleukin 12 (IL-12), potent inducer of IFN-γ.^20^

In cell-to-cell contact-dependent conditions human neutrophils can mimic myeloid-derived suppressor cells and suppress T-cell activation through artefactual mechanisms.^21^ Therefore, CD10^neg^/CD10^pos^ neutrophils were evaluated for their ability to enhance IFN-γ production in cell contact-independent manner. We found that CD10^neg^ neutrophils strongly enhanced IFN-γ and IL-12 production by CD4^+^ T-cells from CMV^+^ patients (Figure 8A, 8B), when co-cultured using a transwell system where CD4^+^ T-cells in the lower chamber were separated from neutrophils in the upper chamber. Of note, CD4^+^ T-cells equally responded to cell-free supernatants derived from CD10^neg^ neutrophils. IFN-γ and IL-12 production were significantly higher in CD4^+^ T-cells from CMV^+^ than CMV^-^ patients. The addition of neutralizing anti-IL-12 antibody abrogated the IFN-γ production by CD4^+^ T-cells from CMV^+^ patients in presence of supernatants derived from CD10^neg^ neutrophils (Figure 8B). These data indicate that CD10^neg^ neutrophils release soluble factors into the culture supernatants that efficiently induce a strong Th1 type response. Further studies aiming at characterizing the neutrophil-secreted immunomodulatory factors are ongoing.

**Figure 8.**
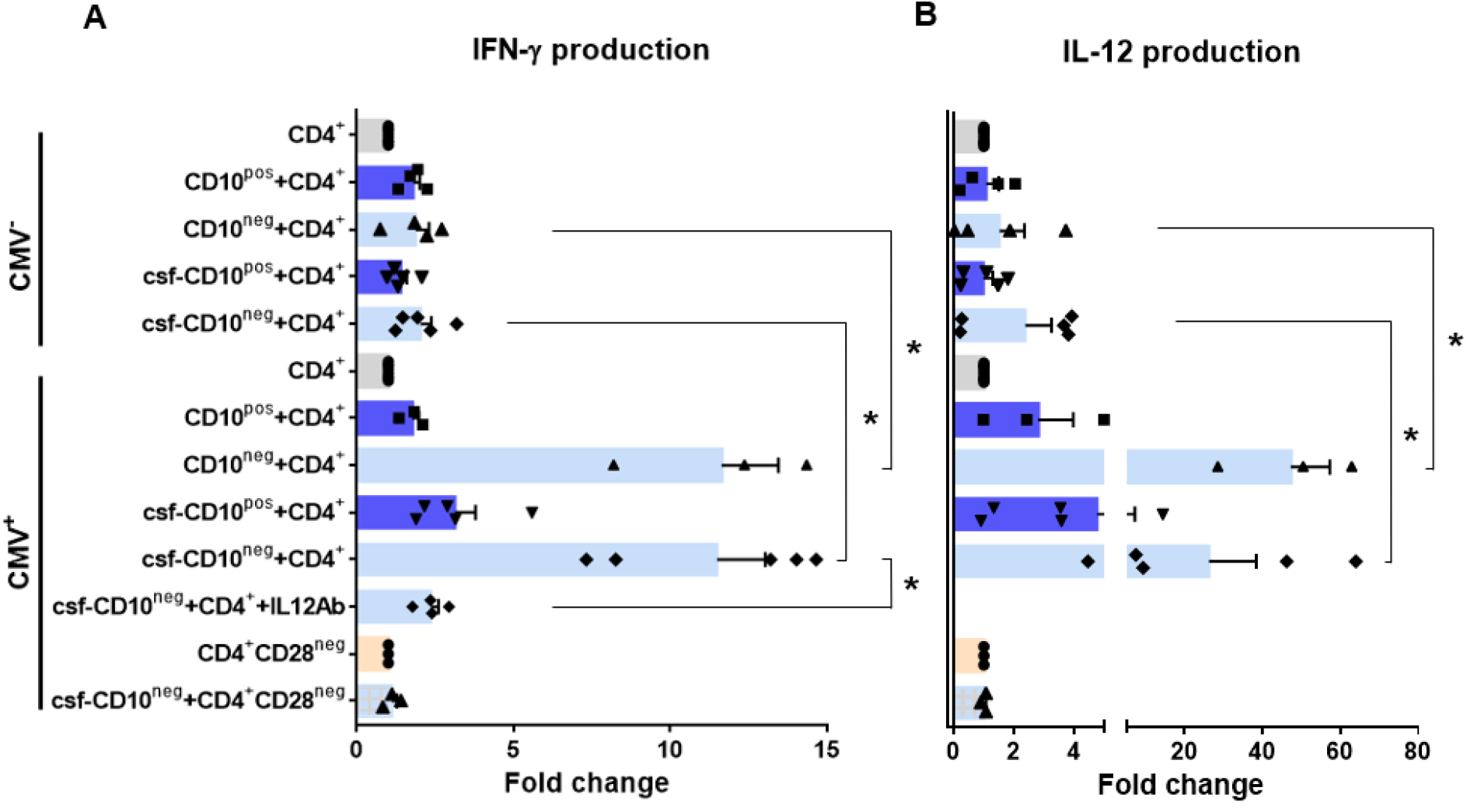
CD10^neg^ neutrophils enhance IFN-γ production by CD4^+^ T-cells via induction of interleukin-12. **A** IFN-γ and **B** interleukin-12 production by CD4^+^ T-cells stimulated with anti-CD3/CD28 beads and co-cultured for 24 hours in absence (CD4^+^) or presence of CD10^pos^ neutrophils (CD10^pos^+CD4^+^), CD10^neg^ neutrophils (CD10^neg^+CD4^+^) using a transwell system or cultured with cell-free supernatants derived from CD10^pos^ neutrophils (csf-CD10^pos^+CD4^+^), CD10^neg^ neutrophils (csf-CD10^neg^+CD4^+^), CD10^neg^ neutrophils in the presence of neutralizing anti-IL-12 antibody (csf-CD10^neg^+CD4^+^+IL12Ab). CD4^+^CD28^null^ T-cells were stimulated with anti-CD3/CD28 beads (CD4^+^CD28^null^) and cultured with cell-free supernatants derived from CD10^neg^ neutrophils (csf-CD10^neg^+CD4^+^CD28^null^). CD10^neg^/CD10^pos^ neutrophils, CD4^+^ T-cells and CD4^+^CD28^null^ T-cells were isolated from CMV-seronegative (CMV^-^) or CMV-seropositive (CMV^+^) patients with AMI (n=3-5). Data are represented as fold-change to respective CD3/CD28 stimulated cells and presented as mean±SEM from independent experiments. \**P*≤0.05.

CD10^neg^ neutrophils had no effect on CD3/CD28-stimulated CD4^+^CD28^null^ T-cells (Figure 8A, 8B), demonstrating that overproduction of IFN-γ is confined to CD4^+^ T-cells expressing CD28. Taken together, our findings indicate that CD4^+^CD28^+^ T-cells from CMV^+^ patients with AMI display a distinct phenotype overproducing IFN-γ in presence of immature neutrophils via induction of interleukin-12.

## DISCUSSION

Innate immune mechanisms play a paramount role during AMI and the functional heterogeneity of monocytes and neutrophils have been the focus of intensive research in recent years. This study highlights for the first time that immature CD16^+^CD66b^+^CD10^neg^ neutrophils and CD14^+^HLA-DR^neg/low^ monocytes promoting proinflammatory immune responses expand in the peripheral blood from patients with large AMI. We also show that immature neutrophils are recruited to the injured myocardium shortly after reperfusion, using a mouse model of AMI. Furthermore, we found a potential link among increased frequency of immature CD10^neg^ neutrophils and elevated IFN-γ levels, especially in cytomegalovirus-seropositive patients with expanded CD4^+^CD28^null^ T-cells. Finally, we could show that CD10^neg^ neutrophils enhance CD4^+^ T-cells IFN-γ production by a contact-independent mechanism involving IL-12.

This study uncovered that CD10 can be used as a surface marker to identify the immature neutrophil population that expands and promotes proinflammatory effects in patients suffering from AMI. We believe that immature CD10^neg^ neutrophils derive from MI-induced emergency granulopoiesis. Both mature (segmented) and immature banded neutrophils are released from the bone marrow presumably to meet the high demand for more neutrophils, especially in patients with large AMI. Not surprisingly, in our study higher frequency of circulating CD10^neg^ neutrophils was associated with increased systemic concentrations of G-CSF, an essential regulator of neutrophil trafficking from the bone marrow to the blood.^15,16^ Recently, CD10 has been proposed as a marker that distinguishes mature from immature neutrophils in healthy volunteers receiving G-CSF for stem cell mobilization.^22^

Multiple clinical trials have evaluated the use of G-CSF in patients with AMI after successful revascularization. The majority of these studies found that effective stem cell mobilization with G-CSF therapy failed to improve left ventricular recovery.^23^ Our findings suggest that the therapeutic benefits of G-CSF therapy after AMI might be compromised due to the release of immature proinflammatory CD10^neg^ neutrophils.

However, neutrophils may be released from the bone marrow in response to increased damage-associated molecular patterns such as S100A8/S100A9, secreted from neutrophils as mediators of sterile inflammation.^24^ Of interest, we found that circulating CD10^neg^ neutrophils express high amounts of S100A9, indicating that immature neutrophils could be an important source of this alarmin in patients with AMI.

Under inflammatory conditions neutrophils traffic to inflamed tissues as well as to draining lymph nodes ^15,25^ modulating T cell-mediated immune responses. Emerging evidence indicates that immature neutrophils can be T-cell suppressive or do possess T-cell stimulatory capacities, displaying disease-specific functional plasticity.^15,26^ Immunostimulatory immature CD10^neg^ neutrophils appear in the circulation of G-CSF–treated healthy volunteers and contact-dependent mechanisms account for their immunoregulatory functions.^22^ Here we provide mechanistic evidence that immature CD10^neg^ neutrophils from patients with AMI, in a contact-independent way involving IL-12, enhance priming for IFN-γ production in activated CD4^+^ T-cells. Thus, through diverse mechanisms immature CD10^neg^ neutrophils may exert immunostimulatory/proinflammatory functions actively participating in the regulation of adaptive immunity.

Genetic and environmental factors shape the immune system over time. Several studies have demonstrated that persistent CMV infection is associated with changes in T-cell phenotype and function.^27–29^ Our results highlight that CD4^+^CD28^+^ T-cells from CMV-seropositive AMI patients are skewed toward a Th1 phenotype, producing large amounts of IFN-γ in presence of CD10^neg^ neutrophils. However, results obtained *in vitro* cannot be translated directly to the *in vivo* situation and several cellular and molecular mechanisms could have led to increased circulating levels of the pleiotropic cytokine IFN-γ after AMI. Notably, using bioinformatic tools (PCA and hierarchical clustering) we were able to highlight the tight relationship among the peripheral expansion of immature CD10^neg^ neutrophils, CMV-altered CD4^+^ T-cell homeostasis and high levels of IFN-γ in patients with large AMI. Thus, determination of circulating CD10^neg^ neutrophils levels, particularly in the context of persistent CMV infection, might help to identify patients at risk for excessive inflammatory immune response.

Although a pathogenetic role of CD4^+^CD28^null^ T-cells in coronary artery disease and atherogenesis have been recognized, important issues have remained unresolved.^30^ A recent study revealed complex associations between of CD4^+^CD28^null^ T-cells and cardiovascular disease.^31^ CD4^+^CD28^null^ T cells are associated with a lower risk for first-time coronary events in a population-based cohort. In contrast, in patients with advanced atherosclerotic disease an increased frequency of CD4^+^CD28^null^ T-cells was associated with more frequent major adverse cardiovascular events. Our findings point to a potential link between CMV induced immune alterations following repeated antigen exposure and the peripheral expansion of CD4^+^CD28^null^ T-cells in ACS patients. CMV has been associated with atherosclerosis and increased risk for cardiovascular diseases. Recent clinical data showed that myocardial ischemia in CMV-seropositive patients leads to significant changes in the composition of the CD8^+^ T-cell repertoire, accelerating immunosenescence.^32^

In spite of numerous studies on polymorphonuclear myeloid cells the presence and functional characteristics of immature neutrophils is underexplored in the setting of AMI in mice. The present study demonstrated for the first time that CD101 can be used as a marker to define the maturation status of neutrophils mobilized into the peripheral blood in response to ischemia and recruited to sites of ischemic injury after reperfusion. Previous studies in a human model of experimental endotoxemia showed that banded neutrophils exhibit efficient migration to sites of infection.^33^ Moreover, developmental analysis of bone marrow neutrophils revealed that immature neutrophils are recruited to the periphery of tumor-bearing mice.^14^ Of note, we found that immature CD101^neg^ neutrophils are released into the bloodstream within minutes after reperfusion and are capable of efficient migration to ischemic tissues, displaying increased expression of MMP-9 and IL-1ß at 3 and 24 hours after reperfusion, respectively. There are significant differences between mouse and human immunology and the transit time of leukocytes may be quite different.^34–36^ During homeostasis, trafficking of neutrophils/myeloid cells from bone marrow into the circulation takes between 1–2 days in mice and 5-8 days in humans.^35^ Such differences should be considered when comparing animal and human studies on immune mechanisms underlying wound healing.

The recruitment of immune cells to sites of tissue repair is a complex highly regulated process involving cytokines, chemokines, and interactions between infiltrating immune cells. HLA-DR^neg/low^ monocytes from patients with AMI are not immunosuppressive but express high amounts of IL1R1. Thus, immature neutrophils, as an important source of IL-1ß in the reperfused heart, may be actively involved in the recruitment of HLA-DR^neg/low^ cells. Saxena *et al*.^37^ showed that IL1R1 signaling mediates early recruitment of Ly6C^hi^ monocytes to the infarcted myocardium. Reperfused myocardial infarction had intense infiltration with Ly6C^hi^ monocytes expressing IL1R1 that peaked after 24 hours of reperfusion.^37^ Noteworthy, recent studies demonstrated that the failing human heart also contains HLA-DR^neg/low^ monocytes.^11^

Several immune mechanisms operate during cardiac wound healing and IFN-γ plays different roles depending on the cellular and microenvironmental context intrinsically linked to the stages of ischemic injury. By integrating cell sorting and *in vitro* experiments we found that macrophages differentiated from HLA-DR^neg/low^ monocytes produced more TNF-α, IL-6, and IL-1ß upon IFN-γ stimulation as HLA-DR^high^ monocyte-derived macrophages. These findings may support a role for HLA-DR^neg/low^ monocytes in pathogenic mechanisms operating during AMI and may, at least in part, explain why an expansion of circulating HLA-DR^neg/low^ monocytes correlates with circulating levels of TNF-α, IL-6, and IL-1ß.

The interleukin-1 pathway plays a key role in post-MI inflammation and the progression to heart failure.^38^ Our *in vitro* mechanistic experiments with immune cells from AMI patients as well as mouse studies provide a potential linkage between the induction of immature CD10^neg^ neutrophils/HLA-DR^neg/low^ monocytes and increased interleukin-1 activity during AMI. Emerging evidences highlight that targeting interleukin-1 may hold promise for patients after MI.^39^ In STEMI patients the interleukin-1 receptor antagonist anakinra significantly reduced the systemic inflammatory response. Moreover, in the CANTOS trial, administration of canakinumab (a monoclonal antibody targeting IL-1β) prevented the recurrence of ischemic events, reduced heart failure-related hospitalizations and mortality in patients with prior AMI.^39^

In conclusion, this study shows that immature CD10^neg^ neutrophils and CD14^+^HLA-DR^lo/neg^ monocytes expand in patients with AMI and highlights their potential role as triggers of immune/inflammatory dysregulation after ischemic injury.

These findings could have major implications for understanding immunoregulatory mechanisms operating during AMI and for the development of future therapeutic strategies. Nevertheless, further studies deciphering the relationship between elevated CD14^+^HLA-DR^lo/neg^ monocytes/CD10^neg^ neutrophils and ensuing mortality and morbidity after ischemic injury are necessary and ongoing.

**Figure 9.**
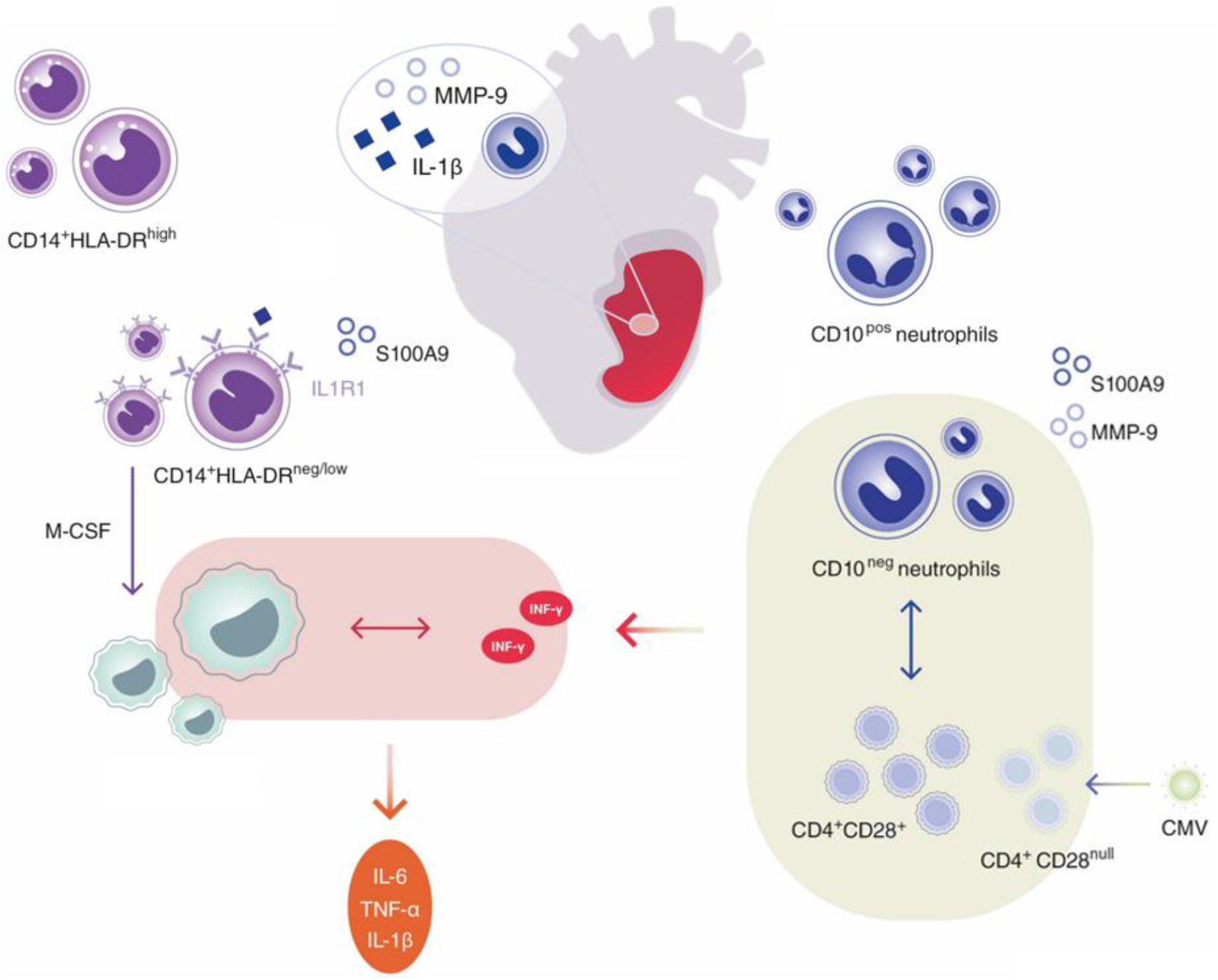
Immature CD10^neg^ neutrophils and HLA-DR^neg/low^ monocytes inducing proinflammatory and adaptive immune responses emerge in patients with large acute myocardial infarction.

## Acknowledgments

We are grateful for the support of Dr Matthias Ballmaier from the Central Research Facility Cell Sorting of the Hannover Medical School.

## Data Availability Statement

All data generated or analysed during this study are included in the manuscript and supporting files.

